# Connexin 36-mediated gap junctions contribute to fine odor discrimination and excitation of mitral cells in the mouse olfactory bulb

**DOI:** 10.1101/2025.09.23.678084

**Authors:** Praveen Kuruppath, Shelly T. Jones, Frederic Pouille, Daniel Ramirez-Gordillo, Diego Restrepo, Nathan E. Schoppa

**Author notes:** Corresponding author: Nathan Schoppa. E-mail addresses. Co-first authors (Praveen Kuruppath, Shelly T. Jones). <u>Author Contributions</u>: All experiments were conducted in the lab of Dr. Nathan Schoppa at UCAMC. PK: Conception and design of study; acquisition, analysis, and interpretation of data; drafting and editing of manuscript. STJ: Conception and design of study; acquisition, analysis, and interpretation of data; drafting and editing of manuscript. FP: Conception and design of study; acquisition, analysis, and interpretation of data. DRG: Acquisition and analysis of data. DR: Acquisition and analysis of data. NES: Conception and design of study; analysis and interpretation of data; drafting and editing of manuscript. All authors approved the final version of the manuscript and agree to be accountable for all aspects of the work in ensuring that questions related to the accuracy or integrity of any part of the work are appropriately investigated and resolved. All persons designated as authors qualify for authorship, and all those who qualify for authorship are listed. <u>Contact for Research Governance at UCAMC</u>: Dr. Thomas Flaig, Vice Chancellor for Research. <u>UCAMC Institutional Animal Care and Use Committee protocol number</u>: 00144.

## Abstract

**Key Points:**

- The output MCs of the olfactory bulb (OB) engage in strong electrical coupling via connexin 36 (Cx36)-mediated gap junctions. However, the behavioral and physiological relevance of these gap junctions is not well understood.
- In studies conducted in Cx36 knock-out (KO) mice, we found that the mice displayed reduced fine odor discrimination capabilities versus wild-type mice in a go/no-go associative learning task. These results provide the first evidence to date of olfactory behavioral deficits in Cx36 KO mice.
- In OB slices, Cx36 KO reduced excitatory responses in MCs to electrical stimulation of sensory afferents, especially during latter stages of the response.
- We suggest that KO-induced impairments in fine odor discrimination are linked to reduced late MC excitation due to the longer time that mice require to make difficult odor discriminations.

The output mitral cells (MCs) and tufted cells (TCs) of the mammalian olfactory bulb (OB) are coupled through both chemical mechanisms as well as gap junctions that are mediated by connexin 36 (Cx36). Here we tested both behavioral and physiological effects of eliminating gap junctions in knockout (KO) mice with homozygous deletions of Cx36. In a go/no-go associative learning task, Cx36 KO mice were found to display reduced discrimination capabilities when presented with pairs of stimuli that included a monomolecular odor and mixtures that had the same monomolecular odor and a small amount of a structurally similar odor. The impairments did not occur for less similar odor pairs, suggesting that Cx36 KO mice have olfactory processing deficits that are specific to fine odor discrimination. In physiological recordings in OB slices from Cx36 KO mice, MCs displayed reduced excitation in response to electrical stimulation of sensory afferents, both single stimulus pulses as well as a theta burst pattern designed to mimic sniffing. The reduction in MC excitatory current was most prominent for late portions of their response, 300 ms after single stimulus pulses or following all theta bursts that came after the first. More global local field potentials recorded in OB glomeruli were largely unaffected by Cx36 KO. We suggest that the KO-induced impairments in fine odor discrimination are linked to reduced late MC excitation due to the longer time that mice require to make difficult odor discriminations.

## Introduction

Studies in recent years have identified multiple mechanisms by which the output cells of the olfactory bulb (OB), the mitral cells and tufted cells (MCs and TCs), can engage in excitatory interactions within glomeruli. Amongst these are chemical mechanisms involving release of the neurotransmitter glutamate from apical dendrites ((Isaacson, 1999; Schoppa and Westbrook, 2002; Urban and Sakmann, 2002; Christie and Westbrook, 2006; De Saint Jan et al., 2009; Najac et al., 2011; Gire et al., 2012; Gire et al., 2019)), along with electrical coupling via gap junctions ((Schoppa and Westbrook, 2002; Hayar et al., 2005; Kosaka and Kosaka, 2005; De Saint Jan et al., 2009; Maher et al., 2009; Gire et al., 2012)). Gap junctional coupling between MCs, which is specific to “sister” MCs that are associated with the same glomerulus, is mediated by connexin (Cx) 36 ((Christie et al., 2005)). Based on studies in Cx36 knock-out (KO) mice, the Cx36-mediated gap junctions appear to function to drive rapidly synchronized spikes in sister MCs ((Christie et al., 2005)). In addition, and perhaps surprisingly, the gap junctions can also contribute to synchronizing non-sister MCs affiliated with different glomeruli ((Pouille et al., 2017)), as part of well-described gamma frequency (40-100 Hz) synchronized oscillations in OB ((Adrian, 1950; Rall and Shepherd, 1968; Kashiwadani et al., 1999)). The gap junction-enhanced “interglomerular” synchrony appears to be at least in part a result of network-level interactions wherein synchronized sister MCs help coordinate GABAergic granule cells (GCs) that are in turn connected to non-sister MCs.

While some of the roles of Cx36-mediated gap junctions in impacting cellular and network properties of the OB are beginning to be understood, there remain no studies to date of the effect of eliminating Cx36-mediated gap junctions on olfactory behavior. Some prior studies that have combined physiological and behavioral measurements *in vivo* have provided some basis for one hypothesis, involving gamma oscillations. For example, during a two-alternate forced choice task, gamma oscillations in the rat OB are strongly associated with more difficult odor discriminations ((Beshel et al., 2007)). Moreover, pharmacological manipulations that alter the magnitude of the oscillations also impair discrimination of odors ((Stopfer et al., 1997; Lepousez et al., 2010; Lepousez and Lledo, 2013)). Thus, reduced gamma oscillations due to Cx36 KO mice ((Pouille et al., 2017)) could result in impaired odor discrimination. Gap junctions could also facilitate odor discrimination through other physiological mechanisms as well, for example if they shape evoked responses that contribute to the tuning of MCs to different odors ((Yokoi et al., 1995; Tan et al., 2010; Gschwend et al., 2016)) or the reliability of neural responsiveness that has been linked to discrimination of sensory stimuli ((Logothetis and Schall, 1989; Britten et al., 1996; Cook and Maunsell, 2002)). There is some evidence from prior recordings in OB slices from Cx36 KO mice that gap junctions contribute to excitation of MCs that is evoked by either stimulation of other MCs ((Christie and Westbrook, 2006)) or olfactory sensory neuron (OSN) afferents ((Vaaga and Westbrook, 2016)).

Here we conducted experiments in wild-type (WT) and Cx36 KO mice to investigate, first, the role of Cx36-mediated gap junctions in modulating olfactory discrimination during a go/no-go task similar to that used in prior studies that showed correlations between gamma oscillations and odor discrimination ((Lepousez and Lledo, 2013)). In addition, we examined the impact of Cx36 KO on excitatory responses in MCs evoked by OSN stimulation in OB slices, expanding on prior studies ((Vaaga and Westbrook, 2016)) by investigating responses under a wide range of stimulus conditions that varied in both intensity and temporal pattern. We also recorded an LFP signal in glomeruli in order to assess the impact of KO on more global excitation in OB that extends beyond MCs. Our results support a role for Cx36-mediated gap junctions in both facilitating odor discrimination and enhancing excitation of MCs, with limited effects on other cell types. We also found that KO effects on MC excitatory responses to OSN stimulation were more complex than previously reported. We propose a model linking the behavioral and physiological results that builds on prior studies that examined the speed-accuracy tradeoff in mice tasked with discriminating odors of differing degrees of similarity ((Abraham et al., 2004; Rinberg et al., 2006; Frederick et al., 2017)).

## Methods

### Ethical Approval

All experiments were approved by the Institutional Animal Care and Use Committee at the University of Colorado Anschutz Medical Campus (UCAMC) in accordance with guidelines set by the U.S. Department of Health and Human Services and outlined in the Public Health Service Policy on Humane Care and Use of Laboratory Animals. The authors understand the ethical principles under which the *Journal of Physiology* operates and our work complies with the *Journal*’s animal ethics checklist.

### Experimental animals

Cx36-LacZ KO mice (Cx36^-/-^, C57BL/6 background), as previously described ((Degen et al., 2004); generous gift of Dr. Richard Benninger at UCAMC), were used for behavioral experiments and the reported patch recordings of MC current and voltage in response to single stimulus pulses applied to OSNs. For local field potential (LFP) and whole-cell patch recordings in response to theta burst stimuli, a different Cx36 KO mouse ((Deans et al., 2001); C57BL/6-129SvEv mixed background) was used; the unpublished data from these mice that are presented here were acquired at the same time as prior recordings of gamma oscillations induced by theta burst stimulation ((Pouille et al., 2017)). While the two Cx36 KO mice were engineered differently and utilized different background mouse strains, they both result in complete elimination of Cx36, as well as similar properties for MC excitatory currents (see below: *Methods: Analysis of electrophysiological recordings*). Behavioral studies were performed in male mice, beginning at postnatal age P35-42. Electrophysiological studies were performed in OB slices prepared from male and female mice at P14–31. Mice were maintained on a 12 h light/dark cycle with food and water ad libitum, except under training and testing conditions for the behavioral experiments (see below). All experimental Cx36 KO (Cx36^-/-^) mice were generated by crossing females and males that were both heterozygous for Cx36 KO. Wild-type (WT) controls for the KO experiments included Cx36^+/+^ littermates as well as unrelated C57BL/6 and C57BL/6-129SvEv mixed strain mice.

### Go/no-go behavioral experiments

In the go/no-go olfactory behavioral experiments, mice were tasked with discriminating an odor that they learned to be associated with a water reward (S+) from an unrewarded odor (S-). Mice were motivated to perform the task by depriving them of water for 1-2 days until they reached 80% of the normal body weight. Behavioral training was performed in the chamber of a computer-controlled olfactometer where they could move freely (Bodyak and Slotnick, 1999; Losacco et al., 2020). Odors were made weekly with high purity odorants (Sigma-Aldrich; vehicle = mineral oil) to a final volume of 10 ml. Volatilized odors (1/40 dilution with air) were presented to mice through an odorant port.

In the go/no-go task, mice self-initiated trials by licking on the water-delivery spout. Licks were detected as electrical connectivity between the waterspout and the ground plate on which they stood. Odors were delivered at a random time 1–1.5 seconds after the first lick. Correct responses occurred when mice licked on the waterspout at least once during each 0.5 second bin in the 2-second response window in order to obtain a water reward for the rewarded S+ odor (a “Hit”) or refrained from licking for the unrewarded S-odor (“Correct rejection”). In trials that were Hits, mice received a ∼10 μl water reinforcement. The mice learned to refrain from licking in response to the S-odor due to the unrewarded effort of sustained licking. Incorrect responses occurred when mice did not lick in response to the S+ odor (a “Miss”) or licked in response to the S-odor (a “False alarm”). Performance was evaluated in blocks of 20 trials, with 10 S+ and 10 S-trials presented at random. Generally, mice underwent 80 trials (four blocks) in a given day. Proficiency in a given block was defined to be a correct response rate of ≥80%, and mice were considered to be proficient in discriminating an odor pair when they displayed proficiency in at least three consecutive blocks on a given day.

All mice first underwent a training phase in which they learned to lick the waterspout to obtain water in the presence of odor (1% isoamyl acetate in mineral oil, v/v) in the ‘begin’ task. Subsequently, they learned to discriminate 1% isoamyl acetate (S+) from mineral oil (S-) in the go/no-go behavior, followed by discrimination of one of the test odors 2-heptanone from mineral oil. If mice reached proficiency in the training tasks, they moved onto the go/no-go testing phase where they discriminated between pairs of odors.

In the testing phase, we assessed the animals’ ability to discriminate between pairs of monomolecular odors as well as between a single monomolecular odor and mixtures that consisted of two odors. We explored two pairs of monomolecular odors with similar molecular structures (2-heptanone versus 3 heptanone and ethyl acetate versus propyl acetate) and associated mixtures as described in the *Results*. For experiments involving mixtures, the rewarded (S+) odors were generally the monomolecular odors (2 heptanone and ethyl acetate) while the unrewarded (S−) odors were the mixtures. The concentrations of the S+ or S-odor stimuli was always 1% in mineral oil. When mixtures were used, the concentrations of the individual components were reduced to bring the total concentration to 1%. To evaluate performance in discriminating odor pairs, we determined the number of days to reach proficiency (for monomolecular odor discriminations) as well as the correct response rates upon reaching proficiency (for monomolecular odor and mixture discriminations). For evaluating the latter parameter, experiments for any given odor pair in which proficient performance was observed were continued for one additional day beyond the day in which proficiency was first shown, such that two days of proficient performance could be evaluated. For the phase of the study involving discrimination of 2-heptanone from 3-heptanone and mixtures of the two, the same sets of seven WT and seven Cx36 KO mice were used for the entire set of odor combinations. For discrimination of ethyl acetate from propyl acetate and mixtures of the two, we used five each of WT and Cx36 KO mice that had been used during the 2-heptanone/3-heptanone discriminations. Two WT and two KO mice dropped out of this phase of the experiments due to lack of motivation.

### Electrophysiological recordings in mouse OB slices

Horizontal slices (300-400 μm) were prepared from OBs of mice following general isoflurane anesthesia and decapitation, as described previously ((Pouille et al., 2017)). Bulb slices were visualized using an upright Axioskop 2FS microscope (Carl Zeiss; Oberkochen, Germany) with differential interference contrast optics video microscopy and a CCD camera. Cells were visualized with a 40X water-immersion objective. All experiments were performed at 29-34°C.

The base extracellular recording solution contained the following (in mM): 125 NaCl, 25 NaHCO_3_, 1.25 NaH_2_PO_4_, 25 glucose, 3 KCl, 2-3 CaCl_2_, 0.5-1 MgCl_2_, pH 7.3, and was oxygenated (95% O2, 5% CO_2_). For whole-cell patch measurements in MCs, identified cells in the MC layer were patched with pipettes (4–8 MΩ) that contained a solution composed of 125 potassium gluconate, 2 MgCl_2_, 0.025 CaCl_2_, 1 EGTA, 2 NaATP, 0.5 NaGTP, 10 Hepes, along with Alexa-488 (50–100 μM) to allow visualization of MC dendritic processes and identification of the MCs’ target glomeruli. The intracellular solution usually also included the sodium channel blocker QX-314 (10 mM) in the patch pipette. For recordings of LFPs in glomeruli, a glass patch-pipette recording electrode filled with extracellular solution (resistance 5–7 MΩ) was placed in the center of an identified glomerulus at the surface of the slice. Whole-cell and LFP recordings were made with a Multi-Clamp 700A or 700B dual patch-clamp amplifier (Molecular Devices, Sunnyvale, CA, USA) and were low-pass filtered at 1–2 kHz using an eight-pole Bessel filter and digitized at 10–15 kHz. Data were acquired using Axograph X software. Whole-cell recordings with a series resistance larger than 20 MΩ were discarded. Epifluorescence was captured via Axiocam HSm (Zeiss) camera; images were acquired using AxioVision software (Zeiss). Reported holding potential (*V*_hold_) values for voltage-clamp experiments were corrected for liquid junction potentials.

OSN stimulation was performed by placing a broken-tip patch pipette (∼10 μm diameter) containing the extracellular solution in the ON layer, ∼50 μm superficial to the target glomerulus of the test MC. The OSN stimulus (100 µs; 10–250 μA intensity), triggered by a biphasic stimulus isolation unit (BSI-950; Dagan, Minneapolis, MN, USA), was applied either as single pulses or as a pattern consisting of four short bursts (five 0.1-ms pulses separated by 10 ms) each in turn separated by 250 ms (4 Hz; theta frequency). Each single stimulus pulse or theta pattern was applied once every 20 s. MCs selected for analysis had apical dendrites targeted to glomeruli near the surface of the OB slice, which facilitated stimulation of OSNs at target glomeruli. MCs were easily identified by their large somas located in the MC layer.

Stimulus-evoked excitatory currents were isolated by recording at *V_hold_* = –77, which would minimize inhibitory currents. In principle, another strategy to eliminate GABAergic inhibition would have been to use a GABA_A_ receptor blocker. However, prior studies have shown that such blockers cause very large increases in prolonged excitatory currents in MCs evoked by OSN stimulation ((Carlson et al., 2000; Schoppa and Westbrook, 2001)). This is because much of the slow current reflects recurrent excitation which is in balance with inhibition. Thus, while blockade of GABA_A_ receptors may have added greater certainty that the recorded currents did not include a contaminating GABAergic current, it would have introduced what we believe is a significantly worse problem in interpreting the slow current and the effect of Cx36 KO on it.

### Analysis of electrophysiological recordings

In the analysis of excitatory currents evoked by OSN stimulation, the peak of the excitatory post-synaptic current (EPSC) reflecting direct monosynaptic input from OSNs (the OSN-EPSC) was defined as the maximum current response within 5 ms of stimulus onset. This window captured the peak of the rapid-onset OSN-EPSC when it was clearly present. For WT, a distinct OSN-EPSC was not always observable at weaker stimulus intensities applied to OSNs even when there was a slower current ((Najac et al., 2011)); in these cases, our method may have captured the start of slower recurrent excitation rather than the OSN-EPSC. Thus, the OSN-EPSCs for WT at weak intensities may have been overestimated. However, this error likely did not significantly impact our conclusion that WT and Cx36 KO had similar OSN-EPSC amplitudes at all OSN stimulation intensities that we systematically examined, including the lowest (50 µA). Even if we assumed that the OSN-EPSC amplitude was zero when WT MCs had no clear OSN-EPSC, we found no significant difference between WT and Cx36 KO for 50 µA stimulation (*p* = 0.21 in Mann Whitney U-test; *n* = 12 MCs for WT, 11 MCs for KO). All reported values for the OSN-EPSC amplitude and other current measurements (see below) reflect absolute magnitudes for the currents.

To evaluate the stimulus strength-dependence of the OSN-EPSCs, the peak OSN-EPSC values were plotted as a function of OSN stimulation intensity and fitted to a sigmoidal function:

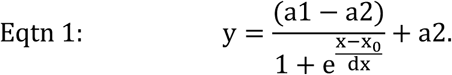

*a2* was defined as the maximum response, *x* was the OSN stimulation intensity, *x_0_* was the stimulus (*stim_1/2_*) that elicited half of the maximum response, *dx* was the steepness of the sigmoidal curve, and *a1* was the minimum response.

In the analysis of the prolonged excitatory current component, estimates of its magnitude were obtained at 30 ms and 300 ms after OSN stimulation. In all MCs from WT mice, the decay of the OSN-EPSC appeared to be complete by 10 ms after the stimulus. Two strategies were used to quantify the magnitude of the prolonged current as a function of the degree of stimulation of OSNs. First, we related the magnitude of the slow current to the absolute current intensity applied to the stimulating electrode. In addition, we related the slow current magnitude to the amplitude of the OSN-EPSC seen in the same data traces. The OSN-EPSCs in these cases provided an estimate of the relative degree of excitation of OSNs at the test MC’s target glomerulus, which in turn contributed to the generation of the slow current. For analysis of MC excitatory currents and glomerular LFPs in response to theta burst stimulation, magnitude measurements were obtained by integrating the current/voltage between the first stimulus pulse and 250 ms after the start of the last stimulus burst (a window of 1 second). Integration windows of 250 ms were used to evaluate the dependence of the MC current response on stimulus burst number. While integrating responses to theta burst stimuli, stimulus artifacts in the current/voltage signals were subtracted out.

As discussed above in the *Experimental Animals* section, we used one type of Cx36 KO mouse ((Degen et al., 2004)) for the analysis of MC currents in response to single OSN stimulus pulses and a second ((Deans et al., 2001)) for responses to theta burst stimulation. We confirmed that the basic observation of the stimulus strength-dependence of the prolonged MC excitatory current that was characterized in KO mice described by Degen and co-workers (see below) also occurred in the other KO mice. In three MC recordings from the mice of Deans and co-workers, the current at 30 ms after the first theta burst stimuli had a small magnitude (mean (SD) = 7 (4) pA) when OSNs were weakly excited (OSN-EPSCs<150 pA) that was similar in value to the 30-ms magnitude observed in the mouse by Degens and co-workers (9 (7) pA for OSN-EPSCs<150 pA, *n* = 9 MCs). Stronger OSN stimulation (OSN-EPSCs>400 pA) recruited much larger currents at 30 ms in both mice (115 (25) pA in mice from Deans and co-workers, *n* = 4 MCs; 176 (81) pA in mice from Degens and co-workers, *n* = 9 MCs).

### Experimental Design and Statistical Methods

All statistical analyses were performed in Prism software (Graph-pad, San Diego, CA). Data are presented as Mean (Standard Deviation). Kruskal-Wallis and Mann-Whitney U statistical tests were used for most comparisons between WT and Cx36 KO. For instances in which multiple comparisons were performed, the Bonferonni correction was applied to the value of the significance level (alpha). Some of the electrophysiological analyses in MCs employed ANCOVA methods to compare the steepness of relationships between different current or voltage parameters and the OSN-EPSC in WT versus Cx36 KO mice. In these cases, we typically pooled measurements obtained at more than one stimulation intensity in a given MC with recordings across all MCs. Similar results were obtained if we used only one measurement per MC, while still sampling a range of OSN-EPSC magnitude values. For example, in the analysis of the MC voltage response (integrated over 0 to 30 ms post-stimulation), we observed a significantly reduced slope value for Cx36 KO (by 59%; *p* = 0.0305) when we used only single measurements from 9 MCs from WT mice and 5 MCs from Cx36 KO mice.

### Data Availability Statement

The data that support the findings of this study are available from the corresponding author upon reasonable request.

## Results

### Cx36 KO mice display deficits in fine odor discrimination

We used the go/no-go odor discrimination task in order to study the role of Cx36-mediated gap junctions on olfactory behavior. In the go/no-go paradigm, mice are tasked with discriminating two odors, one that is associated with a water reward (S+) and a second (S–) that is not (***Fig. 1A***). Mice first underwent a training phase, in which they learned to discriminate isoamyl acetate (1%) as the S+ stimulus from mineral oil (S-) and subsequently one of the test odors 2-heptanone (1%) from mineral oil. During the testing phase, mice were tasked with discriminating various odor pairs (listed in ***Table 1***). Responses were generally evaluated in four 20-trial blocks on a given day, with a response counted as correct either when the mouse licked in response to an S+ odor (“Hit”) or when it did not lick in response to an S-odor (“Correct rejection”). The testing phase generally consisted of examining a mouse’s response to a given odor pair across a sufficient number of days for it reach to proficiency, plus one additional day. Mice were considered proficient during both the training and testing phases when their correct response rate in a 20-trial block was ≥80% across three consecutive blocks. For a few odor pairs in which proficient performance was not observed (see below), testing was stopped after 3 or 4 days.

**Figure 1.**
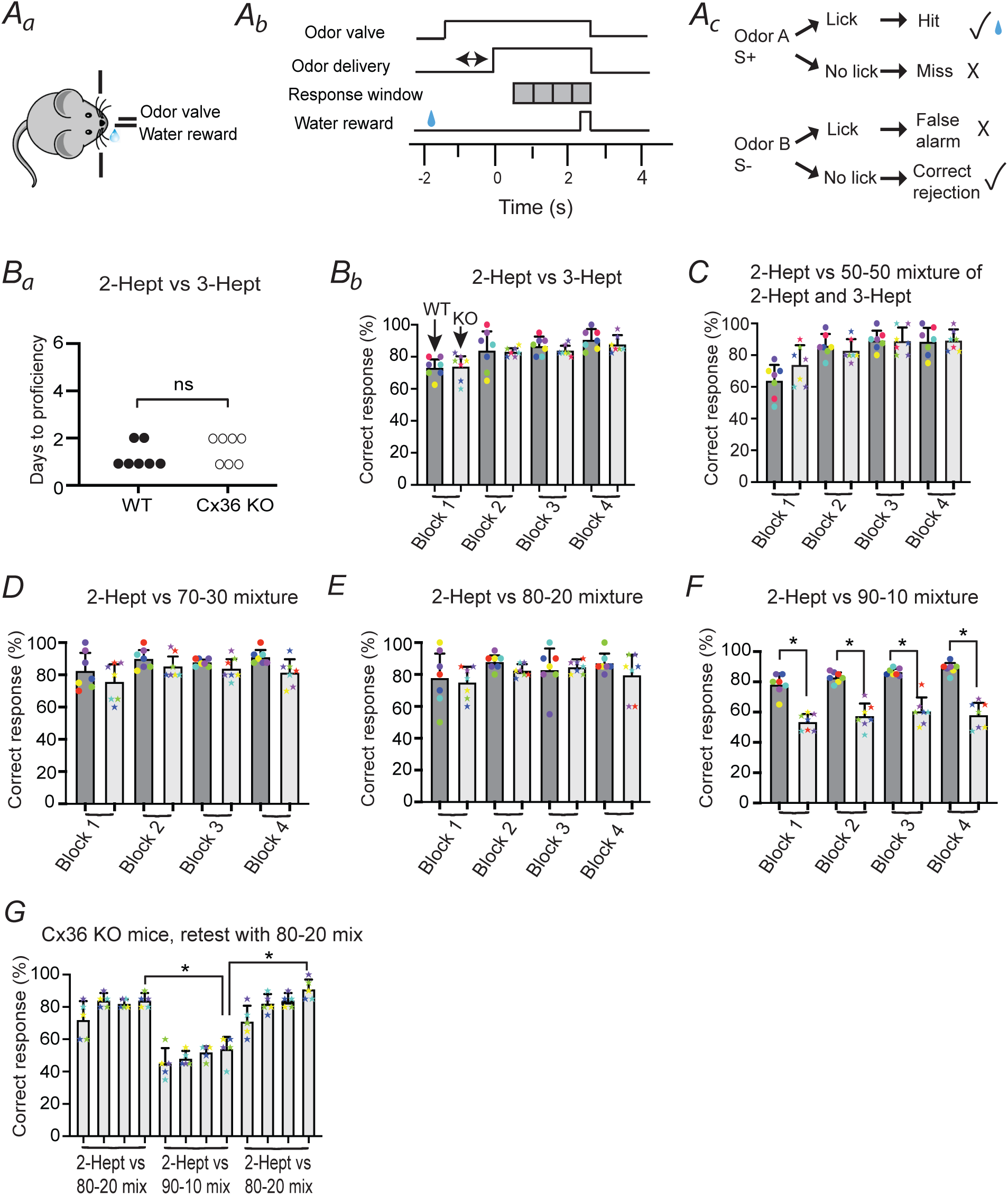
Connexin (Cx) 36 KO mice display deficits in difficult discrimination tasks involving mixtures of heptanones. **(*A*)** Go/no-go olfactory discrimination paradigm. (***A_a_***) Mouse self-initiated trials by licking the water-delivery spout. (***A_b_***) Sequence of steps. Upon the first lick, the odor valve opened with odor delivery being initiated 1-1.5 seconds later and lasting for 2.5 seconds. The last 2 seconds of odor delivery was the response window, when the licking of the mouse was assayed to determine whether it was discriminating a rewarded S+ odor from an unrewarded S– odor. The mouse received a water reward if it licked at least once during each of four 0.5-second blocks of the response window upon delivery of the S+ odor. (***A_c_***) A response was counted as correct (checkmark) if the mouse either licked in response to the S+ odor (Hit) or did not lick in response to S– (Correct rejection). (***B***) *Cx36* KO and WT mice (*n* = 7 for each) were similar at discriminating 2-heptanone (2-Hept, 1%) from 3-heptanone (3-Hept, 1%), as reflected in the fact that they took a similar number of days to reach proficiency (≥80% correct responses; ***B_a_***) and displayed similar correct response rates upon reaching proficiency (***B_b_***). Data in ***B_b_*** and in parts ***C***-***F*** reflect correct response rates for each of four consecutive blocks on the last two of 2-4 days of testing. Error bars reflect standard deviations (SDs). The correct response rates for individual mice are superimposed on the bar graphs, color-coded for each mouse. (***C***-***F***) Mixture experiments in which mice were discriminating 2-Hept from mixtures that included 2-Hept and 3-Hept at varying contributions between 50% 2-Hept/50% 3-Hept (50-50; ***C***), 70-30 (***D***), 80-20 **(*E***), and 90-10 (***F***). Note that KO mice performed as well as WT mice in all cases except for discrimination involving the most difficult 90-10 mixture. \**p* < 0.001 in Mann-Whitney U-test. (***G***) Cx36 KO mice that failed to discriminate 2-Hept from the 90-10 2-Hept/3-Hept mixture were successful at discriminating the easier 80-20 mixture upon retesting. Data reflect five KO mice that were subject to retesting, showing performance for the 80-20 mixture prior to the 90-10 mixture (left-most bars), for the 90-10 mixture (middle bars), and during the first day of re-exposure to the 80-20 mixture. The data points for the first 80-20 and the 90-10 exposures reflect the mean performances for the last two days of testing under those conditions for these five mice. \**p* = 0.0079 in Mann-Whitney U-test comparing the fourth blocks under each condition.

**Table 1.**
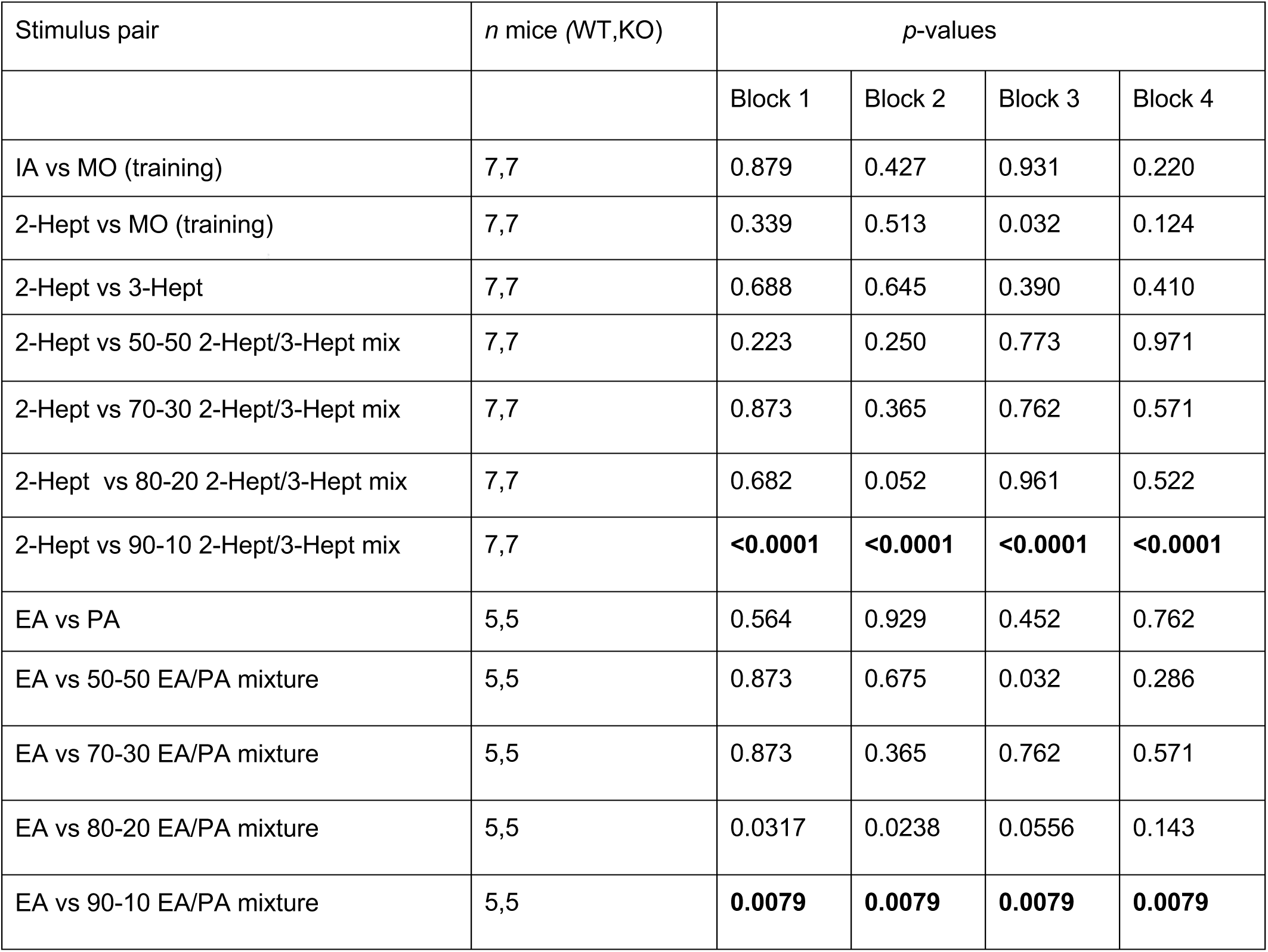
Summary of stimulus pairs used and *p*-values in go/no-go odor discrimination experiments. Results of Mann Whitney U-tests applied to values of correct response rates for WT and Cx36 KO mice across four 20-trial blocks. Values used were the mean correct response rates observed for each mouse during the last two days of experiments for each stimulus pair. Applying the Bonferroni correction for multiple (4) comparisons, *p* < 0.0125 was considered to be significant (indicated in bold). IA = isoamylacetate; MO = mineral oil; 2-Hept = 2-heptanone; 3-Hept = 3-heptanone; EA = ethyl acetate; PA = propyl acetate.

| Stimulus pair | *n* mice (WT,KO) | *p*-values |  |  |  |
| --- | --- | --- | --- | --- | --- |
|  |  | Block 1 | Block 2 | Block 3 | Block 4 |
| IA vs MO (training) | 7,7 | 0.879 | 0.427 | 0.931 | 0.220 |
| 2-Hept vs MO (training) | 7,7 | 0.339 | 0.513 | 0.032 | 0.124 |
| 2-Hept vs 3-Hept | 7,7 | 0.688 | 0.645 | 0.390 | 0.410 |
| 2-Hept vs 50-50 2-Hept/3-Hept mix | 7,7 | 0.223 | 0.250 | 0.773 | 0.971 |
| 2-Hept vs 70-30 2-Hept/3-Hept mix | 7,7 | 0.873 | 0.365 | 0.762 | 0.571 |
| 2-Hept vs 80-20 2-Hept/3-Hept mix | 7,7 | 0.682 | 0.052 | 0.961 | 0.522 |
| 2-Hept vs 90-10 2-Hept/3-Hept mix | 7,7 | **<0.0001** | **<0.0001** | **<0.0001** | **<0.0001** |
| EA vs PA | 5,5 | 0.564 | 0.929 | 0.452 | 0.762 |
| EA vs 50-50 EA/PA mixture | 5,5 | 0.873 | 0.675 | 0.032 | 0.286 |
| EA vs 70-30 EA/PA mixture | 5,5 | 0.873 | 0.365 | 0.762 | 0.571 |
| EA vs 80-20 EA/PA mixture | 5,5 | 0.0317 | 0.0238 | 0.0556 | 0.143 |
| EA vs 90-10 EA/PA mixture | 5,5 | **0.0079** | **0.0079** | **0.0079** | **0.0079** |

In the testing phase, we first compared the ability of WT and Cx36 KO mice to discriminate structurally similar monomolecular odors, 2-heptanone (2-Hept, 1%; S+) and 3-heptanone (3-Hept, 1%; S-). For this odor pair, we found that WT and KO mice displayed similar levels of discrimination, as reflected in the number of days to reach proficiency (***Fig. 1B_a_***; WT mean (SD):1.28 (0.45) days, *n* = 7 mice; Cx36 KO: 1.57 (0.49) days, *n* = 7 mice; *p* = 0.592 in Mann-Whitney U-test) and performance upon reaching proficiency (***Fig. 1B_b_***). When we analyzed the combined results from the last two days of testing, no significant differences were observed between WT and Cx36 KO mice in any of the blocks (***Table 1***). Next, the odor discrimination task was made progressively more difficult by having mice discriminate 2-Hept (S+) from mixtures of 2-Hept and 3-Hept (S-), wherein the contribution of 2-Hept in the mixture increased across each stage. Mixtures included 2-Hept and 3-Hept in ratios of 50%-50% (50-50), 70-30, 80-20, and 90-10, while keeping the total odor concentration at 1%. When the mixtures were 50-50, 70-30, or 80-20, no differences between WT and KO mice were observed (***Fig. 1C-E***, ***Table 1***). However, for the most difficult discrimination task involving the 90-10 mixture of 2-Hept and 3-Hept, performance in Cx36 KO mice was markedly worse than in WT mice (***Fig. 1F***, ***Table 1***), with KO mice performing at close to chance level (50% correct responses). The poorer performance of Cx36 KO mice in the 90-10 mixture experiments occurred in spite of the fact that KO mice were generally given one additional day of testing versus WT mice (2.26 (0.71) days for WT, *n* = 7; 3.14 (0.34) days for Cx36 KO, *n* = 7; *p* = 0.0047 in Mann-Whitney U-test). These results imply that Cx36 KO mice can perform quite well in discriminating similar odors, as do WT mice, but they show impairments at the most difficult discriminations.

The structure of the discrimination experiments, wherein the task became progressively more difficult across stages, raised the possibility that the reduced performance for the 90-10 2-Hept/3-Hept mixture was because KO mice were more susceptible than WT mice to fatigue or loss of interest in the discrimination task. To verify that this was not the case, we retested the ability of five of the Cx36 KO mice that had just failed to discriminate the 90-10 mixture by re-exposure to the 80-20 mixture on the subsequent day. We found that these mice were quickly able to reach proficiency in discriminating the 80-20 mixtures (***Fig. 1G***). By the last block on the first day of retesting, the mice were much better at discriminating the 80-20 mixture (91 (6)% correct responses) than during the last block of the prior two days of experiments with the 90-10 mixture (54 (8)% correct responses; *p* = 0.0079 in Mann-Whitney U-test, *n* = 5).

After the WT and Cx36 KO underwent go/no-go experiments with 2-Hept and 3-Hept and their associated mixtures, a subset of them (five of each mouse type) were subject to discrimination tasks involving structurally similar acetates, ethyl acetate (EA) and propyl acetate (PA). As with the individual heptanones, WT and Cx36 KO mice performed similarly when discriminating EA (1%, S+) from PA (1%, S-) both in terms of number of days to reach proficiency (***Fig. 2A_a_***, 1.20 (0.40) days for WT, *n* = 5 mice; 2.40 (2.80) days for Cx36 KO, *n* = 5 mice; *p* >0.999 in Mann-Whitney U test) and correct response rates upon reaching proficiency (***Fig. 2A_b_***, ***Table 1***). When discriminating EA from mixtures of EA and PA at different ratios (50-50 EA-PA, 70-30, 80-20, and 90-10), KO mice performed as well as WT mice for the easier 50-50 and 70-30 mixtures (***Fig. 2B,C***, ***Table 1***) but displayed markedly impaired performance for the most difficult discrimination involving the 90-10 mixture as compared to WT mice (***Fig. 2E***, ***Table 1***). For the somewhat easier task involving discrimination of EA from the 80-20 EA-PA mixture, KO mice did not display a significant difference in performance when data were analyzed in a block-by-block fashion (***Fig. 2D***, ***Table 1***), but differences were observed when all blocks were combined for each mouse (87 (6)% correct responses for WT, *n* = 5 mice; 71 (6)% correct responses for Cx36 KO, *n* = 5 mice; *p* = 0.0159). Thus, overall, as in the experiments with heptanones, Cx36 KO mice displayed specific impairments in discrimination of EA from EA-PA mixtures with only the most difficult mixtures.

**Figure 2.**
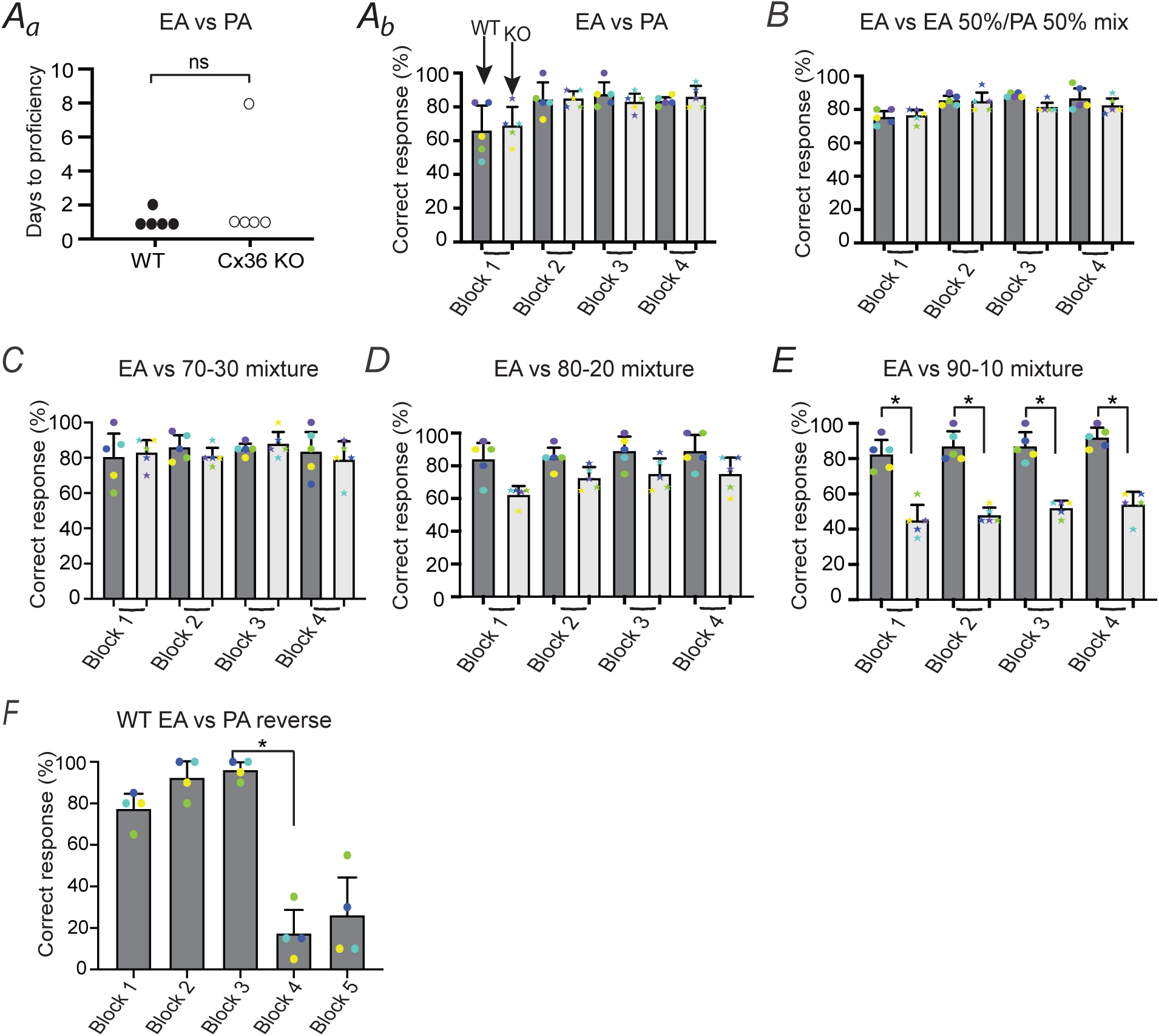
Cx36 KO mice display deficits in more difficult discrimination tasks involving mixtures of acetates. (***A***) Cx36 KO and WT mice were similar at discriminating ethyl acetate (EA, 1%) and propyl acetate (PA, 1%), as reflected in the number of days to reach proficiency (≥80% correct responses; ***A_a_***) and correct response rates upon reaching proficiency (***A_b_***). Data in ***A_b_*** and in parts ***B***-***E*** reflect correct response rates for each of four consecutive blocks on the last two days of 2-9 days of testing. The color coding for individual mice was the same as in Fig. 1. (***B***-***E***) Mixture experiments in which mice were discriminating EA from mixtures that included EA and PA at varying contributions including 50% EA/50% PA (50-50; ***B***), 70-30 (***C***), 80-20 **(*D***), and 90-10 (***E***). Note that KO mice performed as well as WT mice in all cases except for the most difficult 90-10 mixture. \**p* = 0.0079 in Mann-Whitney U-test, *n* = 5 for each mouse type. (***F***) For EA versus PA discrimination, an additional day of testing was added for WT mice wherein the S+ and S– odors were reversed after the third of five blocks (PA = S+, EA = S-following reversal). Reversal resulted in a large decrease in correct response rate. \**p* = 0.0286 in Mann-Whitney U-test, *n* = 4.

In the odor discriminations that involved the individual odors EA and PA (not mixtures), we added an additional day of testing for WT mice in which we increased the number of 20-trial blocks from four to five and reversed the identity of the S+ and S– odors between the third and fourth blocks (S+ = PA, S– = EA). We found that the correct response rate decreased substantially, from 96 (4)% for Block 3 to 18 (10)% for Block 4 (***Fig. 2F***; *p* = 0.0286 in Mann-Whitney U-test, *n* = 4). This argued that the test animals were not relying on other cues associated with the S+ and S-stimuli, for example auditory cues derived from opening of the water reward valve when they were successful at discriminating.

A final issue that we addressed in the behavioral experiments was whether the impaired odor discrimination in Cx36 KO mice for the most difficult mixtures reflected deficits in sensory processing versus learning deficits. The fact that discrimination deficits in KO mice were specific to the most difficult mixtures by itself suggested a mechanism involving impaired sensory processing, but we also turned to data obtained during the training phase of the go/no-go experiments to evaluate whether Cx36 KO mice had olfactory learning deficits. When mice were discriminating the first training odor isoamyl acetate from mineral oil, WT and KO mice (*n* = 7 for each) took a similar number of days to reach proficiency (***Fig. 3A_a_***; 4.29 (0.88) days for WT, 5.14 (1.23) days for KO, *p* = 0.217 in Mann-Whitney U-test) and also displayed similar correct response rates upon reaching proficiency (***Fig. 3A_b_***, ***Table 1*).** WT and Cx36 KO mice also performed similarly when discriminating 2-Hept from mineral oil in terms of days to reach proficiency (***Fig. 3B_a_***; 4.00 (2.13) days for WT, 5.14 (0.98) days for KO, *p* = 0.188 in Mann Whitney U test) and correct response rates (***Fig. 3B_b_***, ***Table 1***). These results suggest that Cx36 KO mice have normal olfactory learning capabilities.

**Figure 3.**
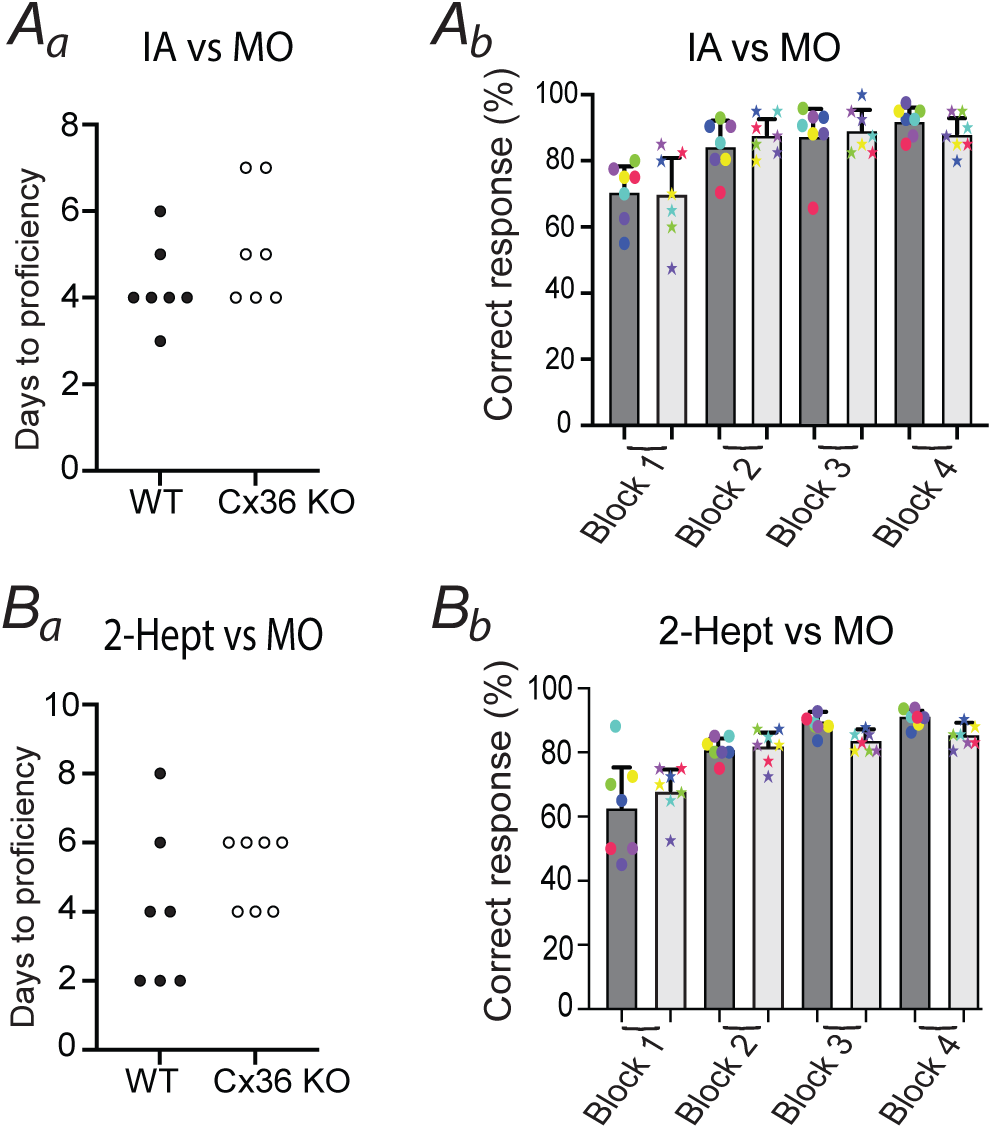
Evidence that impaired odor discrimination in Cx36 KO mice was not due to learning deficits. (***A***) In the training phase involving discrimination of isoamyl acetate (IA, 1%) from mineral oil (MO), Cx36 KO mice performed similarly as WT mice, taking a similar number of days to reach proficiency (***A_a_***) and displaying similar correct response rates upon reaching proficiency (***A_b_***). (***B***) In the training phase involving discrimination of 2-Hept from MO, Cx36 KO mice also took a similar number of days to reach proficiency as WT mice (***B_a_***) and displayed similar correct response rates upon reaching proficiency (***B_b_***). Data in ***A_b_*** and ***B_b_*** reflect the last two days of experiments in the IA vs MO and 2-Hept vs MO discriminations.

### Cx36 KO has complex attenuating effects on MC excitation

Given our behavioral results suggesting that Cx36 KO impairs fine odor discrimination, we next assessed the impact of Cx36 KO on cellular and circuit properties of the olfactory bulb (OB), which is the primary olfactory processing structure widely believed to facilitate odor discrimination ((Friedrich and Wiechert, 2014; Mori and Sakano, 2021)). Prior studies in OB slices have already provided evidence for one circuit-level effect of Cx36 KO on the bulb involving a reduction in gamma oscillations ((Pouille et al., 2017)). Our focus here was mainly on the magnitude of excitatory responses of output mitral cells (MCs) to electrical stimulation of olfactory sensory neurons (OSNs) at varying levels of intensity (100 µsec, 10-200 µA; ***Fig. 4A***). While electrical coupling via gap junctions is not restricted to MCs, also extending to TCs and GABAergic neurons (Hayar et al., 2005; Kosaka et al., 2005), coupling appears to be particularly strong for MCs ((Gire et al., 2012)) and thus their responses were more likely to be impacted by Cx36 KO.

**Figure 4.**
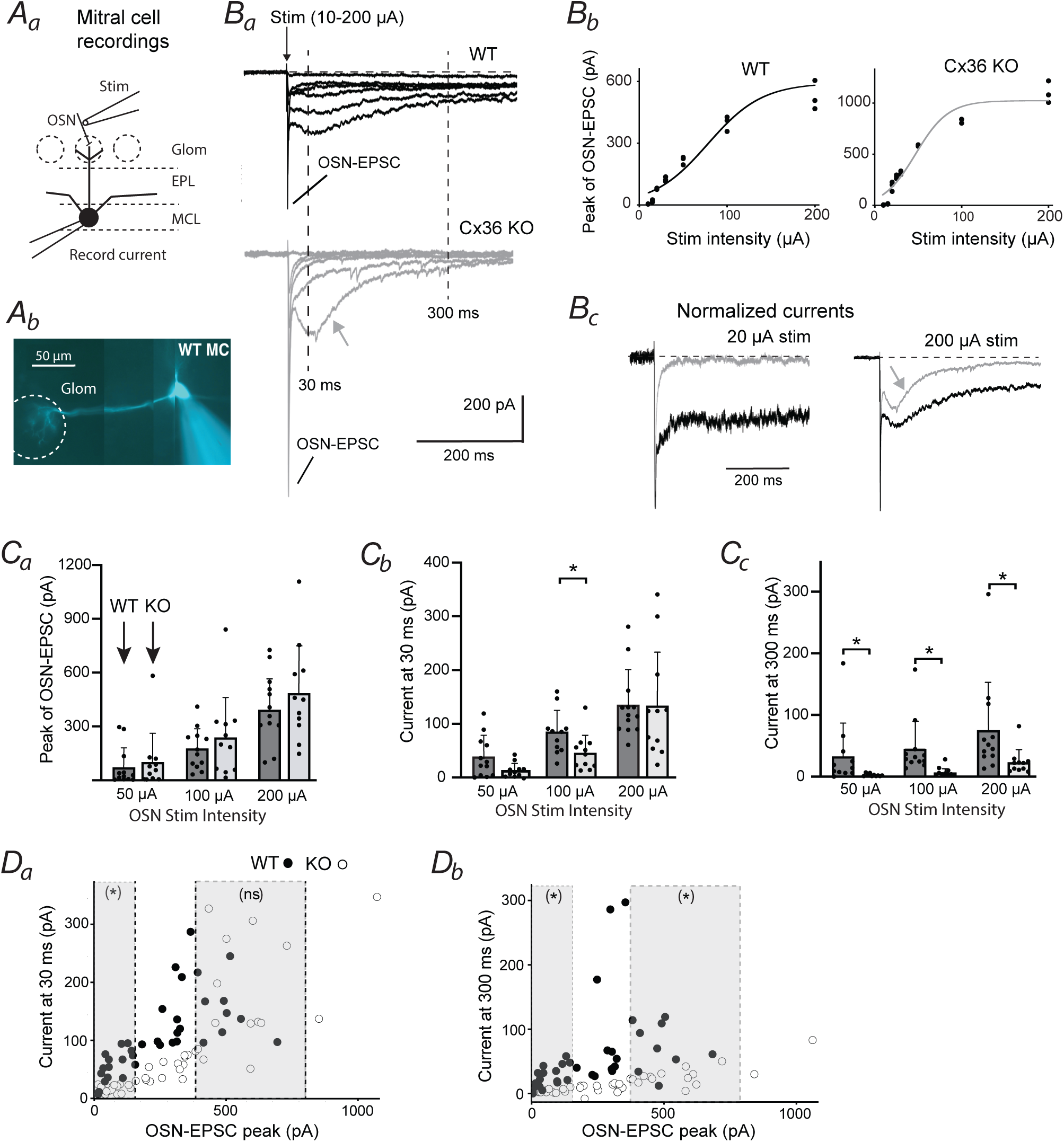
Cx36 KO has complex attenuating effects on mitral cell (MC) excitatory currents. (***A***) General protocol. (***A_a_***) Excitatory currents in a MC were recorded (at *V_hold_* = –77 mV) in response to single stimulus pulses applied to olfactory sensory neurons (OSNs) at the MC’s target glomerulus. (***A_b_***)The MC’s target glomerulus was deduced by following a MC’s Alexa-488-filled apical dendrite. Glom = glomerulus, EPL = external plexiform layer, MCL = mitral cell layer. (***B***) Example current recordings in MCs in OB slices prepared from WT and Cx36 KO mice. (***B_a_***) Currents recorded across a wide range of OSN stimulation intensities (10, 20, 30, 50, 100, 200 µA) in WT (top, black traces) and KO mice (bottom, gray traces). The currents in MCs from both WT and KO mice included both a short-duration monosynaptic OSN-EPSC (evident as a fast “spike” of current at this time-scale) along with a much longer-lasting current. Note that the MC from KO mice displayed a prominent ∼100 ms-duration current with strong 200 µA stimuli (gray arrow) that is qualitatively similar to the current in the WT MC. Data traces reflect averages of 3 trials. (***B_b_***) Plots relating the peak amplitude of the OSN-EPSCs (absolute magnitudes) to OSN stimulation intensity were fitted to sigmoidal function (see *Methods*) in order to estimate the mid-point voltage of activation of the OSN-EPSC. In these recordings, the mid-point voltages were 78 µA for WT and 47 µA for KO. (***B_c_***) MC currents from part ***B_a_*** at weak (20 µA) and strong (200 µA) OSN stimulation intensities that were normalized to the peak of the OSN-EPSC. Note that at 20 µA, the KO MC displayed essentially no prolonged current, but such a current emerged at 200 µA (gray arrow). (***C***) Summary of current measurements sorted by the absolute intensity of OSN stimulation. Shown are magnitude measurements for the OSN-EPSC (***C_a_***), as well as the current measured at 30 ms (***C_b_***) and 300 ms (***C_c_***) time-points after OSN stimulation. \**p* ≤ 0.0091. Measurements reflect recordings from the same 12 WT MCs and 11 Cx36 KO MCs. (***D***) Summary of current measurements (absolute magnitudes) at 30 and 300 ms time-points (from parts ***C_b_*** and ***C_c_***) sorted by the relative degree of OSN activation, as estimated from the magnitude of the OSN-EPSCs (from part ***C_a_***). (***D_a_***) At 30 ms, the MC current was much smaller in KO mice versus WT when the OSN-EPSCs were <150 pA (data points in left shaded region; \**p* = 0.0006, Mann-Whitney U-test) but similar in size for OSN-EPSCs>400 pA (right shaded region). (***D_b_***) For MC currents at the 300 ms time-point, KO caused a reduction regardless of OSN-EPSC amplitude. \**p* ≤ 0.0068 in Mann-Whitney U-test.

As previously reported ((Najac et al., 2011; Vaaga and Westbrook, 2016; Jones et al., 2020)), single stimulus pulses applied to OSNs typically evoked excitatory currents in MCs in WT mice (recorded at *V_hold_* = –77 mV) with a complex waveform that included a fast-onset monosynaptic OSN-EPSC along with a prolonged current that lasted many hundreds of milliseconds (***Fig. 4B_a_***). Evaluating, first, the effect of Cx36 KO on the OSN-EPSC, we found no significant differences between KO and WT (***Fig. 4C_a_***, ***Table 2*)**. This was true across a wide range of stimulation intensities systematically tested (50, 100, 200 µA) that resulted in a ∼10-fold difference in the average magnitude of the OSN-EPSC. Cx36 KO also did not impact the stimulus-dependence of the OSN-EPSC. In fitted sigmoidal curves to the stimulus-response relationships for the OSN-EPSCs for WT and KO (examples in ***Fig. 4B_b_***), the stimulus intensity values that led to half-maximal OSN-EPSCs were similar between WT and KO (80 (15) μA for WT, *n* = 10; 105 (42) μA for Cx36 KO, *n* = 9; *p* = 0.278, Mann-Whitney U-test). It should be noted that the OSN-EPSC magnitudes in MCs from both WT and Cx36 KO mice were highly variable at any given stimulus intensity across recordings (***Fig. 4C_a_***), which may have obscured small effects of KO on OSN-EPSCs. This variability may have reflected differences in the structural organization of OSN fibers at different glomeruli on the surface of the OB slices. Some of the MCs’ target glomeruli had clearly identifiable bundles of OSN fibers on which stimulating electrodes could be placed, leading to large stimulation-evoked MC currents even at weak intensities, but many target glomeruli did not display such bundles at the slice surface.

**Table 2.**
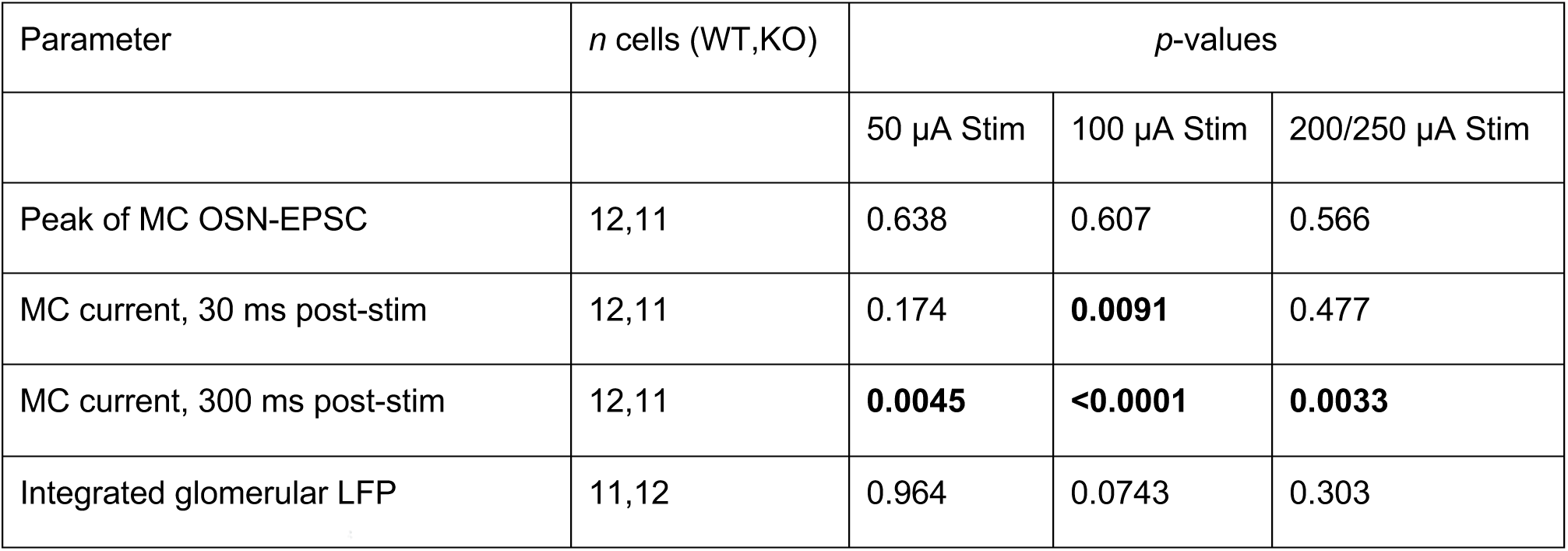
Statistical comparisons of electrophysiological parameters between WT and Cx36 KO mice. Results of Mann-Whitney U-tests applied to electrophysiological parameters for WT and Cx36 KO mice across three OSN stimulation intensities. The first three parameters were derived from current recordings from mitral cells (MCs), the last from measurements of LFPs in glomeruli. The highest intensity was 200 µA for MC current recordings and 250 µA for LFPs. Applying the Bonferroni correction for multiple (3) comparisons, *p* < 0.0167 was considered to be significant (indicated in bold).

While Cx36 KO did not significantly impact the monosynaptic OSN-EPSC in MCs, KO had clear effects on more prolonged currents. Indeed, at weaker stimulation intensities, MCs in Cx36 KO mice sometimes displayed essentially no discernible prolonged current (see normalized traces at 20 µA in ***Fig. 4B_c_***). We found however that KO did not simply eliminate the current that followed the OSN-EPSC ((Vaaga and Westbrook, 2016)), as more complex effects were revealed when we analyzed the MC current at different OSN stimulation intensities and also paid attention to different time points after OSN stimulation. As can be seen in both the example raw and normalized traces in ***Figure 4B_a_*** and ***4B_c_***, stronger stimulation of OSNs resulted in a current in MCs from KO mice that lasted ∼100 ms. At the strongest stimulation intensity consistently tested (200 µA), this current in KO mice was just as large as in WT, as assessed from the current measured at the 30-ms time-point after OSN stimulation (***Fig. 4C_b_***; 135 (63) pA for WT, *n* = 12; 133 (99) pA for KO, *n* = 11; *p* = 0.477 in Mann-Whitney U-test). That KO shifted the dependence of the ∼100 ms current on the level of OSN activation was also supported by an additional analysis (***Fig. 4D_a_***) in which, across many recordings, we related the magnitude of the current at the 30-ms time point in a given MC and stimulation intensity (values in ***Fig. 4C_b_***) to the OSN-EPSC that was observed in the same traces (values in ***Fig. 4C_a_***). Here the OSN-EPSC magnitude was used as an estimate of the relative degree of OSN activation at the MCs’ target glomeruli; this strategy at least partially circumvented the problem of variable efficacies of stimulating OSN axons across MC recordings (see above). This analysis showed that for smaller OSN-EPSCs that were less than 150 pA, the current in WT (43 (29) pA; 18 measurements at different stimulus intensities in 9 cells) was ∼4-fold larger than in KO mice (9.3 (7.0) pA; 19 measurements at different stimulus intensities in 9 cells; *p* = 0.0006 in Mann-Whitney U-test), but this difference disappeared when the OSN-EPSCs were greater than 400 pA (WT: 139 (49) pA, 8 measurements at different stimulus intensities from 6 cells; Cx36 KO: 176 (93), 11 measurements at different stimulus intensities from 9 cells; *p* = 0.585 in Mann-Whitney U test).

The complex effect of KO on the current lasting ∼100 ms however did not extend to currents measured at much longer times after OSN stimulation. We found that, at the 300-ms time point after OSN stimulation, the MC current was consistently reduced in Cx36 KO mice by 70-90% regardless of whether the data were sorted by OSN stimulation intensity (***Fig. 4C_c_***, ***Table 2***) or the magnitude of the OSN-EPSC (***Fig. 4D_b_***; *p* < 0.0001 in Mann-Whitney U-test for OSN-EPSCs < 150 pA; *p* = 0.0068 for OSN-EPSCs > 400 pA; *n* values are the same as in analysis conducted above on the 30 ms-current). That KO may have different effects on MC excitatory currents at different time-points is consistent with the more prolonged current that extends beyond the OSN-EPSC reflecting multiple mechanisms (see *Discussion*). Taking the results of the measurements at 30 ms and 300 ms together, the effects of Cx36 KO can be summarized as a reduction in the excitatory current in MCs that followed the OSN-EPSC, although the precise effect at different time-points depended on the degree of activation of OSNs.

What effect does Cx36 KO have on the MC voltage response at different levels of OSN activation? We found, similar to previous reports ((Christie et al., 2005)), that eliminating gap junctions in KO mice caused a ∼60% increase in the input resistance of MCs (79 (23) MΩ for WT, *n* = 12; 126 (23) MΩ for KO, *n* = 11; *p*<0.0001, Mann-Whitney U-test; ***Fig. 5A***), as would be expected if KO eliminated a significant membrane conductance. Thus, it was possible that the generally smaller excitatory currents in MCs in KO mice could still result in relatively large depolarizations. To examine this issue, we maintained an approach in which we used the OSN-EPSC as a measure of the relative level of OSN activation, and related the MC voltage response in current-clamp to the OSN-EPSC recorded in voltage-clamp in the same cell and stimulation intensity (***Fig. 5B***).

**Figure 5.**
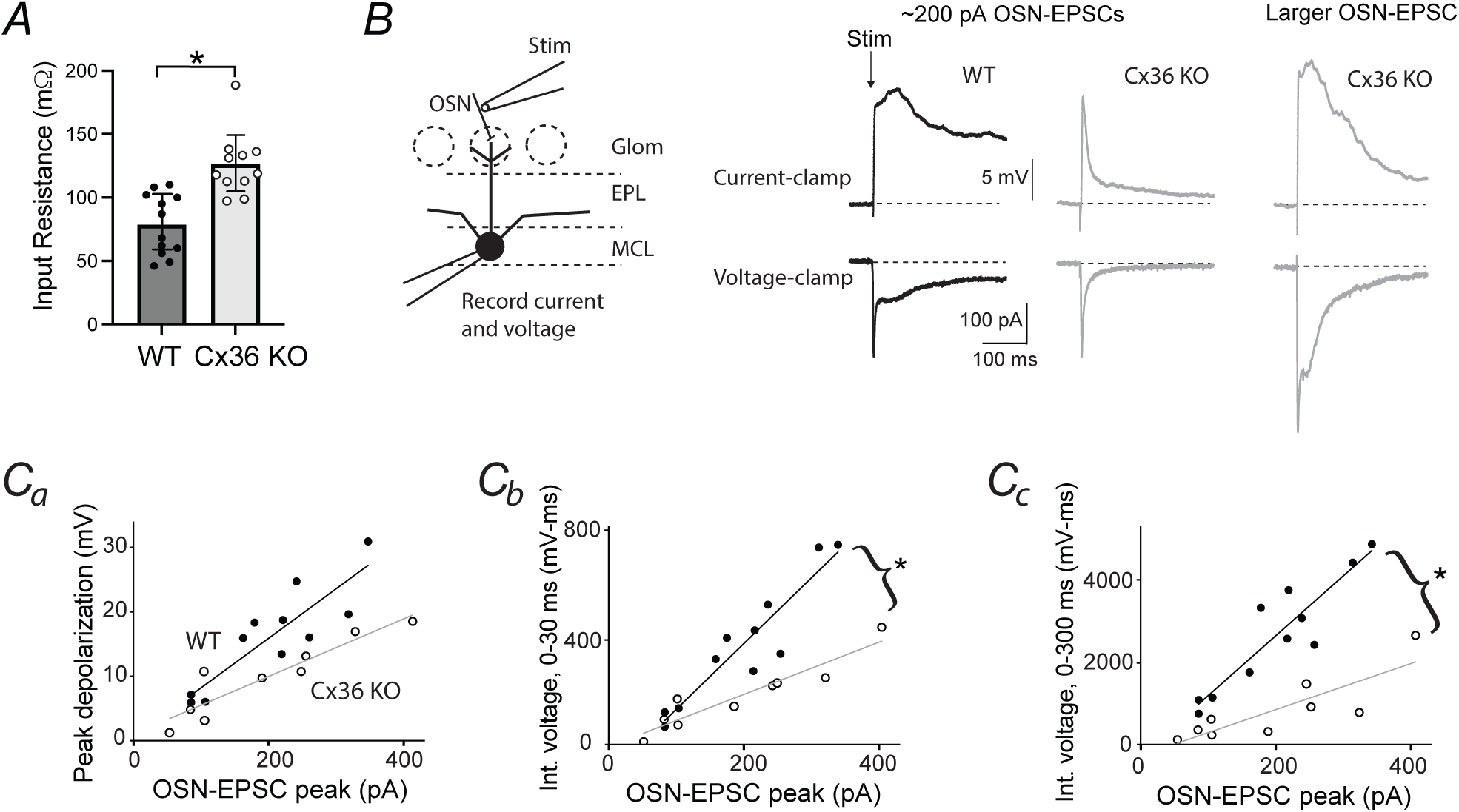
Cx36 KO reduces MC voltage responses to OSN stimulation. (***A***) Input resistance measurements for MCs in WT (*n* = 12) and Cx36 KO (*n* = 11) mice. \**p* < 0.0001 in Mann-Whitney U-test. (***B***) Example recordings of voltage and current responses in the same MCs. The traces at left (WT) and middle (KO) were chosen such that the OSN-EPSCs had similar magnitudes of ∼200 pA, i.e., there was a similar level of OSN activation. Note the much longer-lasting, large voltage response in MCs from WT versus KO mice, reflecting the fact that the underlying current in the same MCs had a delayed component in WT that was mainly absent in KO. The traces at right show that the voltage response in the same KO MC did become longer-lasting at higher levels of OSN activation (larger OSN-EPSCs) when there was a prolonged excitatory current. The recordings included the sodium channel blocker QX-314 (10 mM) in the patch pipette, which allowed us to assess the effect of KO on voltage deflections more easily without contamination from over-riding action potentials. (***C***) Relationship between relative OSN activity levels, as estimated from the magnitude of the OSN-EPSC, and the MC voltage response for WT and KO mice. Three different measures of the voltage response are shown, including peak depolarization (***C_a_***), integrated voltage, 0-30 ms post-stimulation (***C_b_***), and integrated voltage, 0-300 ms post-stimulation (***C_c_***). Asterisks next to the plots in ***C_b_*** and ***C_c_*** indicate significantly reduced slopes for KO (*p* ≤ 0.0035), as determined by ANCOVA analysis. The 11 plotted values for WT in each of the plots were taken from 9 MC recordings in which OSN stimulation intensity was varied; the 9 plotted values for KO were taken from 5 MC recordings.

Furthermore, we used ANCOVA methods to assess the effect of KO on MC voltage responses across different OSN-EPSC magnitudes (***Fig. 5C***), where any effect of KO would be seen as a difference in the slope of lines fitted to plots relating MC voltage to the OSN-EPSC. This showed that Cx36 KO reduced the evoked voltage response of MCs when it was assessed by integrating the voltage response across either the first 30 ms (slope reduced by 60%; *p* < 0.001; 11 measurements at different stimulus intensities in 9 MCs for WT, 9 measurements at different stimulus intensities in 5 MCs for KO) or the first 300 ms (slope reduced by 64%; *p* = 0.0035) after OSN stimulation. Thus, in spite of KO increasing MC input resistance, KO reduced the evoked MC depolarization, reflecting the smaller prolonged components of the excitatory current. It should be noted that by one other measure of the strength of the MC depolarization, the peak value, KO had more inconsistent effects (slope reduced by 42%; *p* = 0.0561). This likely reflected the fact that the peak depolarization was sometimes shaped by the magnitude of the monosynaptic OSN input, which was not different between KO and WT (***Fig. 4C_a_***).

As one other test of the effect of Cx36 KO on MC excitation, we recorded MC currents in response to theta burst stimulation applied to OSNs ((Hayar et al., 2004; Pouille et al., 2017); four bursts of five 0.1-ms pulses, each separated by 250 ms, 4 Hz; ***Fig. 6A***). Theta burst stimulation was designed to mimic the sniffing cycle-dependent activation of OSNs seen under physiological conditions ((Kay, 2014)). The currents were quantified by integrating the current over a 1-second window that included the four stimulus bursts, a measure that should have captured the composite of KO effects on the different current components identified in responses to single OSN stimulation pulses (***Fig. 4C***). As above, we related the integrated current to the magnitude of the OSN-EPSC as an estimate of degree of OSN activation (***Fig. 6B***), using the OSN-EPSC evoked by the first stimulus in the stimulus train (see inset for KO traces in ***Fig. 6A***). This analysis indicated that Cx36 KO caused a 69% reduction in the slope of lines relating the integrated excitatory current to the OSN-EPSC (*p* = 0.0004 in ANCOVA analysis; WT: 16 measurements at different stimulation intensities from 8 MCs for WT; 16 measurements at different stimulation intensities from 8 MCs for KO). Thus, the KO-induced reduction in excitatory responses in MCs extends to conditions in which physiological-like stimuli are applied.

**Figure 6.**
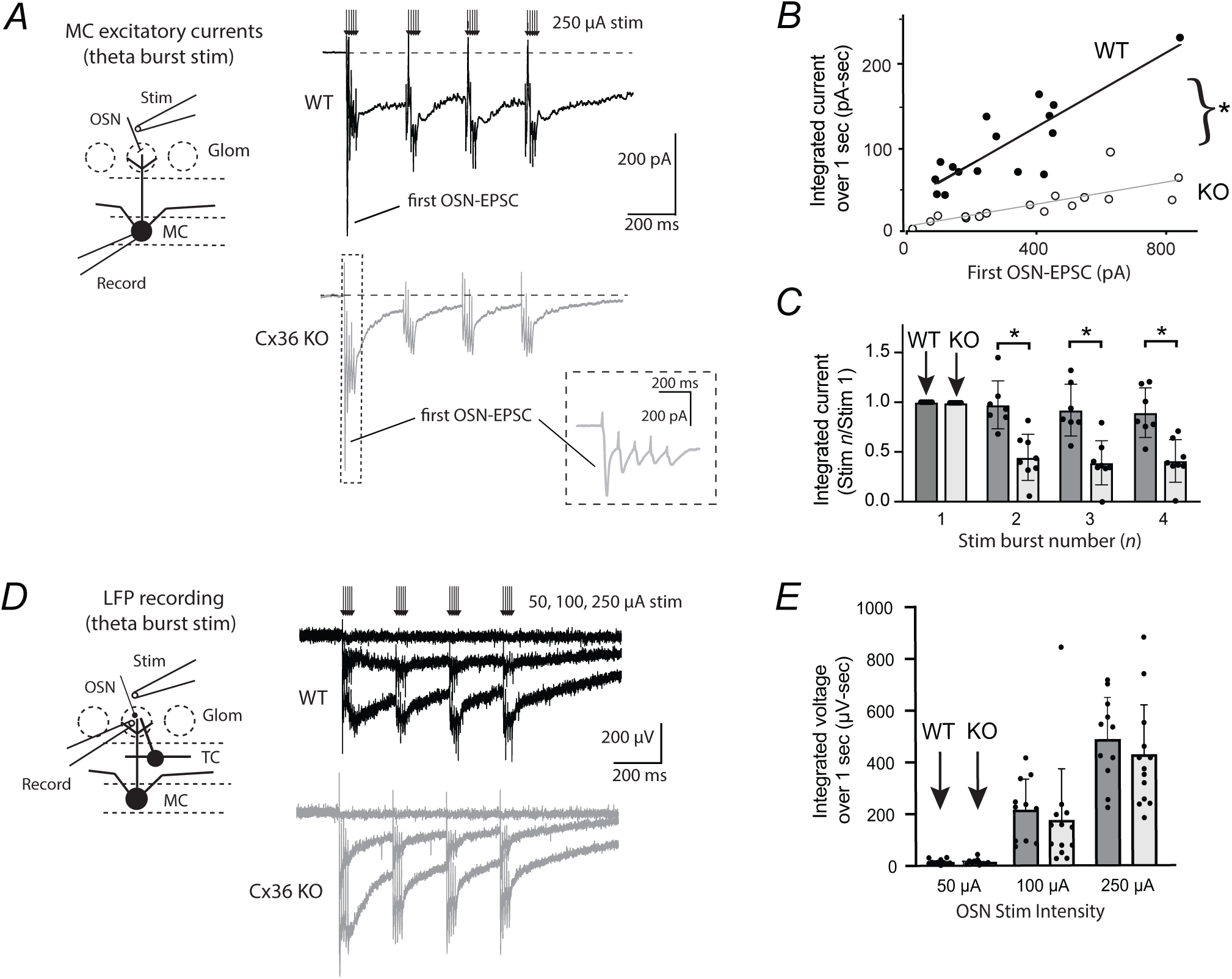
During theta burst stimulation of OSNs, Cx36 KO reduces MC excitatory currents but not glomerular local field potentials (LFPs). (*A*) Example recordings of MC excitatory currents (at *V_hold_* = −77 mV) in response to theta burst stimulation of OSNs (250 µA) in WT (top) and Cx36 KO mice (bottom). Inset shows expansion of the MC current response in KO mice in response to the first burst of OSN stimuli, illustrating depressing monosynaptic OSN-EPSCs ((Vaaga and Westbrook, 2016)). The magnitude of the first OSN-EPSC in this burst was used to estimate the relative degree of OSN activation in part *B*. (*B*) Summary of MC current measurements in response to theta burst stimulation, as reflected in plot of OSN-EPSC magnitude versus integrated current (over 1 sec). The observed steeper slope for WT (\**p* = 0.0004 in ANCOVA analysis) supports that MCs in WT mice had a stronger excitatory response to theta burst stimulation. The 16 plotted values for WT were from 8 MC recordings in which OSN stimulation intensity was varied; the 16 values for KO were from 8 MC recordings. (*C*) Across the four bursts of stimuli, MCs from Cx36 KO mice displayed a stronger reduction in evoked current (integrated over 250 ms) between the first and all subsequent bursts. \**p* < 0.0012 in Mann-Whitney U-test, *n* = 7 WT MCs, *n* = 8 KO MCs. (*D*) Example recordings of LFPs recorded in glomeruli in response to theta burst stimulation in WT (top) and Cx36 KO mice (bottom). LFPs from both mouse types are shown in response to 50, 100, and 250 µA stimulation. Except at the lowest stimulation intensity, the LFPs in both had a significant long-lasting component after each stimulus burst. (*E*) Summary of integrated LFP measurements over 1 second at different intensities: 50, 100, and 250 µA. No significant differences were observed between WT (*n* = 11) and KO (*n* = 12) at any of the intensities.

The MC current recordings conducted with theta burst stimulation revealed one other interesting aspect of the MC response to repeated stimulation: it appeared that the attenuating effect of Cx36 KO on MC excitatory currents was not uniform across the sniff-like pattern, being more pronounced for currents that appeared after the second, third, and fourth stimulus bursts versus the first burst (see data traces in ***Fig. 6A***). This was also apparent in an analysis in which we normalized the current measured after each stimulus burst (integrated across 250 ms) to that elicited by the first burst (***Fig. 6C***). The currents associated with the second, third, and fourth stimulus bursts in Cx36 KO were about 50% of the corresponding relative currents in WT (e.g., ratio of second versus first burst response: 0.97 (0.24) for WT, *n* = 7 MCs; 0.45 (0.23) for KO, *n* = 8 MCs; *p* = 0.0006 from Mann-Whitney U-test). Thus, MCs in Cx36 KO mice not only displayed reduced excitatory responses during theta burst stimulation but were also not able to sustain them as well beyond the first stimulus burst.

### Lack of effect of Cx36 KO on the glomerular LFP

In the final part of the study, we sought to assess the effect of Cx36 KO on OB circuit properties that extend beyond the responsiveness of MCs. Here, rather than systematically evaluating all of the different individual cell-types, including the various subtypes of TCs ((Imamura et al., 2020)), we evaluated the effect of Cx36 KO on a more global measure of the responsiveness of the bulb to OSN stimulation: the local field potential (LFP) that can be recorded in glomeruli (***Fig. 6D***; (Gire and Schoppa, 2009; Pouille et al., 2017)). This “glomerular LFP” mainly reflects signals originating from the apical dendrites of MCs and TCs, in addition to processes of some other cell-types that reside at glomeruli. We examined KO effects on LFP responses to theta burst stimulation of OSNs in order to enable comparisons to KO effects on MC excitatory currents recorded under the same condition (***Fig. 6A***). Interestingly, we found that Cx36 KO had no observable effect on the glomerular LFP across a range of OSN stimulation intensities (***Fig. 6E***, ***Table 2***), in contrast to the strong effect that KO had on MC excitatory currents.

That Cx36 KO had at least much smaller effects on glomerular LFP versus the MC response was not surprising as MCs engage in much stronger electrical coupling versus other cell-types in which this has been quantitatively assessed (i.e., external tufted cells; Gire et al., 2012). On the other hand, one might have expected KO to have some effect on the glomerular LFP, as it does partially reflect signals originating at MC apical dendrites. Possible explanations for the absence of an apparent effect of KO on the LFP include the fact that other cell-types, including TCs, significantly outnumber MCs ((Shepherd, 2004)). Also, potential small effects of KO on the glomerular LFP may have been obscured by the relatively high degree of variability in LFP response magnitude across recordings at a given OSN stimulation intensity (***Fig. 6E***). This variability likely reflected the same issues in the organization of OSN axons at glomeruli at the OB slice surface that contributed to the high degree of variability of MC current amplitudes at given OSN stimulation intensities (**Fig. 4C**; see above).

## Discussion

In this study, we sought to examine behavioral and physiological deficits in the olfactory system in Cx36 KO mice, building on prior observations that MCs in OB that are associated with the same glomerulus engage in strong electrical coupling through Cx36-mediated gap junctions ((Christie et al., 2005)). We found that eliminating the gap junctions in Cx36 KO mice caused impairments in fine discrimination between odors. Cx36 KO also reduced excitation of MCs in responses to sensory input, although more global excitatory responses in OB were not significantly impacted. We discuss these results below, including how they may be related to each other.

### Behavioral deficits in odor discrimination in Cx36 KO mice

Our evidence that Cx36 KO impairs fine odor discrimination was based on experiments involving a go/no-go associative learning task in which mice had to distinguish different pairs of odors in order to receive a water reward. The odor pairs tested included different but structurally similar monomolecular odors (2-Hept versus 3-Hept and EA versus PA) as well as pairs that included one of the monomolecular odors and mixtures of differing ratios that included the same monomolecular plus a second, structurally similar monomolecular odor. We found that both WT and Cx36 KO mice were in fact quite proficient at discriminating similar odor pairs, but KO mice had deficits for the most difficult discrimination tasks involving mixtures (e.g., 2-Hept versus a 2-Hept/3-Hept mixture with 90% 2-Hept). These results suggest that Cx36-mediated gap junctions contribute to the processing of information about odors that facilitate odor discrimination but mainly only under conditions in which the discrimination task is quite difficult.

We considered the possibility that reasons other than sensory processing deficits could have contributed to the impaired discrimination performance in Cx36 KO mice. For example, the design of the behavioral experiments was such that the discrimination abilities for any given mouse was tested on pairs of odors that became progressively more similar, raising the possibility that the discrimination failure for KO mice for the most difficult tasks was due to fatigue or loss of interest. However, we controlled for this by showing that KO mice that had displayed impaired performance for the most difficult discrimination could still succeed when tested immediately afterwards with an easier discrimination (***Fig. 1G***). It was also unlikely that our results could have been explained by memory impairments ((Frisch et al., 2005; Allen et al., 2011)) or heightened anxiety ((Zlomuzica et al., 2012)) in Cx36 KO mice. If they had been, the deficits in KO mice wouId likely have extended to more difficult odor discriminations. We also directly evaluated KO effects on learning/memory based on the performance of Cx36 KO mice during the training phase of the study (***Fig. 3***). Here we found that KO and WT mice took the same number of days to reach proficiency and displayed similar proficiencies.

To our knowledge, our results reflect the first studies to date that have examined the behavioral consequences of Cx36 KO on olfaction. Nevertheless, there were some caveats. Amongst these is the fact that our experiments were conducted in mice in which Cx36 was eliminated throughout development of the mice. Thus, there could have been compensatory changes in the OB circuit to account for loss of Cx36-mediated gap junctions. If so, it is possible that our studies may have underestimated the role of gap junctions; perhaps the actual function of gap junctions in facilitating odor discrimination could extend to somewhat easier discriminations. We also did not explicitly test for KO effects on the sensitivity of the mice to odors, which is an analysis that is generally more difficult to perform in mice versus evaluating discrimination between odors. Given our physiological results suggesting a KO-induced reduction in MC excitation (see below), we would predict that Cx36 KO mice would have reduced sensitivity.

### Physiological underpinnings for the effect of Cx36 KO on olfactory discrimination

In the electrophysiological analysis that we conducted in parallel with the behavioral experiments, we mainly examined MC excitatory responses to stimulation of OSNs, either single stimulus pulses or a theta burst pattern that mimicked sniffing. In the responses to single stimulus pulses, Cx36 KO had complex effects that depended on the OSN stimulation intensity and also what part of the excitatory current response examined (***Fig. 4C***). The most dramatic effect of KO was on a prolonged current component that occurred hundreds of milliseconds after OSN stimulation. KO reduced the current recorded at the 300 ms time point after stimulation by 70-90% across all OSN stimulation intensities. In contrast KO had little to no effect on the rapid-onset monosynaptic EPSC reflecting direct inputs from OSNs ((Vaaga and Westbrook, 2016)) at all OSN stimulation intensities. Cx36 KO had an effect that was, in a sense, between these two on an intermediate-duration current that lasted ∼100 ms after OSN stimulation. KO did not impact the magnitude of this current at high OSN stimulation intensities but did shift the range of OSN activity required to recruit the current. These effects of Cx36 KO on MC currents are more complex than a prior report that suggested that KO essentially eliminated the current that followed the OSN-EPSC ((Vaaga and Westbrook, 2016)); however, the prior study did not examine responses to a range of OSN stimulation intensities. During responses to theta-burst stimulation, Cx36 KO reduced the integrated evoked current in MCs by quite a large amount, about 70%. However, KO effects during theta burst stimulation were not uniform across the train, being larger on MC currents evoked by latter stimulus bursts that followed the first (***Fig. 6C***). This suggests that, in a Cx36 KO mouse engaged in sampling an odor with sniffing, MCs would be less able to sustain an excitatory response.

How might these attenuating effects of Cx36 KO on MC excitation be related to reduced fine olfactory discrimination capabilities that we observed during go/no-go experiments? One framework for considering our results is provided by prior behavioral studies that utilized the go/no-go olfactory discrimination paradigm to evaluate the time it takes for mice to accurately discriminate odors of different degrees of similarity ((Abraham et al., 2004)). These investigators found that highly accurate discrimination between monomolecular odors required about 200 ms, but this time increased to 300-400 ms for more difficult tasks involving odor mixtures. That accurate discrimination of odors requires more time for more similar odor pairs has also been shown using other experimental paradigms besides go/no-go as well ((Rinberg et al., 2006; Frederick et al., 2017); but see (Uchida and Mainen, 2003)). In the context of these behavioral studies, our electrophysiological results showing that Cx36 KO had larger effects on later components of the MC current response are notable, both the especially large effect of KO on the single stimulus-evoked MC current at the 300 ms time point as well as the poor ability of MCs in KO mice to sustain an excitatory current during theta burst stimulation. We would speculate (***Fig. 7***) that the effect of KO on difficult odor discriminations reflects the large KO-induced reduction in the MC current at the critical later time points when information is still being gathered for difficult behavioral decisions. A Cx36 KO-induced reduction on MC excitation at later time points could impact behavioral discrimination capabilities through changes in response reliability ((Logothetis and Schall, 1989; Britten et al., 1996; Cook and Maunsell, 2002)) or a reduction in gamma frequency (40-100 Hz) synchronized oscillations in OB that have been implicated in fine odor discrimination ((Stopfer et al., 1997; Beshel et al., 2007)). Gamma oscillations, which involve the back-and-forth interplay between MCs and GABAergic granule cells ((Rall and Shepherd, 1968; Lagier et al., 2004; Galan et al., 2006; Schoppa, 2006a, b)), have already been shown to be reduced in Cx36 KO mice ((Pouille et al., 2017)).

**Figure 7.**
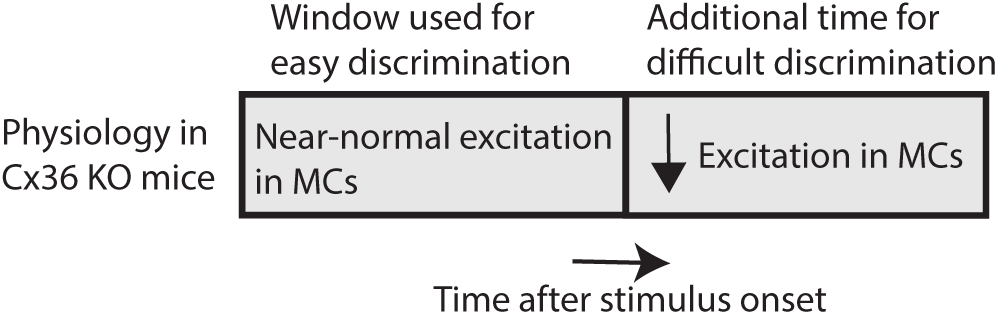
Working mechanistic model for impaired fine odor discrimination in Cx36 KO mice. The amount of time it takes for accurate behavioral performance depends on the difficulty of the discrimination task, with difficult tasks involving similar odor pairs requiring more time after stimulus onset ((Abraham et al., 2004; Rinberg et al., 2006; Frederick et al., 2017)). MCs in KO mice have relatively normal levels of excitation during the early time-window when easy discriminations can be made but much reduced excitation during later times when information is still being integrated for difficult discriminations.

A final point of interest, more at the level of cellular mechanisms, is what might account for the variety of different effects that Cx36 KO had on MC excitatory responses to OSN stimulation. One explanation for the shift in the OSN activity level required to recruit the intermediate (∼100 ms)-duration current component was provided by the prior study by Vaaga and Westbrook (2016). They reported that Cx36 KO, while not impacting the monosynaptic OSN-EPSC in MCs, did cause a moderate reduction in the monosynaptic EPSC in eTCs. Because eTCs can mediate feedforward excitation between OSNs and MCs (De Saint Jan et al., 2009; Najac et al., 2011; Gire et al., 2012) a KO-induced reduction in monosynaptic OSN signals in eTCs could mean that higher levels of OSN activity would be necessary to drive the same level of eTC-driven feedforward excitation. Also fitting with this explanation for the shift in the dependence of the ∼100 ms-duration current on OSN activation is the fact that, in eTC-MC pair-cell recordings, the main part of the MC current response to bursts of action potentials in eTCs lasts about 100 ms ((De Saint Jan et al., 2009)). As for the largest effect of Cx36 KO, on the most prolonged current, the 70-90% reduction would be consistent with most of the current in a given MC being driven by ion channel mechanisms located on other MCs to which it is electrically-coupled rather than on that MC itself. The most likely candidate mechanism for the prolonged current involves activation of metabotropic glutamate receptor 1 ((Schoppa and Westbrook, 2001; Heinbockel et al., 2004; Ennis et al., 2006; Yuan and Knopfel, 2006; De Saint Jan and Westbrook, 2007; Vaaga and Westbrook, 2017)). Certainly, many mechanistic questions remain, including why the drastic effect of Cx36 KO was limited to the most prolonged current. The varying effects of Cx36 KO on the different MC current components might reflect differences in the localization of the relevant synapses and glutamate receptors versus the gap junctions ((Rash et al., 2005; Bourne and Schoppa, 2017)) or perhaps varying effects of developmental compensation in the Cx36 KO animal.

## Acknowledgements

We thank members of the Schoppa lab for helpful discussions.

## References

Abraham NM, Spors H, Carleton A, Margrie TW, Kuner T, Schaefer AT (2004) Maintaining accuracy at the expense of speed: stimulus similarity defines odor discrimination time in mice. Neuron 44:865–876.

Adrian ED (1950) The electrical activity of the mammalian olfactory bulb. Electroencephalography and clinical neurophysiology 2:377–388.

Allen K, Fuchs EC, Jaschonek H, Bannerman DM, Monyer H (2011) Gap junctions between interneurons are required for normal spatial coding in the hippocampus and short-term spatial memory. The Journal of neuroscience: the official journal of the Society for Neuroscience 31:6542–6552.

Beshel J, Kopell N, Kay LM (2007) Olfactory bulb gamma oscillations are enhanced with task demands. The Journal of neuroscience: the official journal of the Society for Neuroscience 27:8358–8365.

Bodyak N, Slotnick B (1999) Performance of mice in an automated olfactometer: odor detection, discrimination and odor memory. Chemical senses 24:637–645.

Bourne JN, Schoppa NE (2017) Three-dimensional synaptic analyses of mitral cell and external tufted cell dendrites in rat olfactory bulb glomeruli. The Journal of comparative neurology 525:592–609.

Britten KH, Newsome WT, Shadlen MN, Celebrini S, Movshon JA (1996) A relationship between behavioral choice and the visual responses of neurons in macaque MT. Vis Neurosci 13:87–100.

Carlson GC, Shipley MT, Keller A (2000) Long-lasting depolarizations in mitral cells of the rat olfactory bulb. The Journal of neuroscience: the official journal of the Society for Neuroscience 20:2011–2021.

Christie JM, Westbrook GL (2006) Lateral excitation within the olfactory bulb. The Journal of neuroscience: the official journal of the Society for Neuroscience 26:2269–2277.

Christie JM, Bark C, Hormuzdi SG, Helbig I, Monyer H, Westbrook GL (2005) Connexin36 mediates spike synchrony in olfactory bulb glomeruli. Neuron 46:761–772.

Cook EP, Maunsell JH (2002) Dynamics of neuronal responses in macaque MT and VIP during motion detection. Nature neuroscience 5:985–994.

De Saint Jan D, Westbrook GL (2007) Disynaptic amplification of metabotropic glutamate receptor 1 responses in the olfactory bulb. The Journal of neuroscience: the official journal of the Society for Neuroscience 27:132–140.

De Saint Jan D, Hirnet D, Westbrook GL, Charpak S (2009) External tufted cells drive the output of olfactory bulb glomeruli. The Journal of neuroscience: the official journal of the Society for Neuroscience 29:2043–2052.

Deans MR, Gibson JR, Sellitto C, Connors BW, Paul DL (2001) Synchronous activity of inhibitory networks in neocortex requires electrical synapses containing connexin36. Neuron 31:477–485.

Degen J, Meier C, Van Der Giessen RS, Sohl G, Petrasch-Parwez E, Urschel S, Dermietzel R, Schilling K, De Zeeuw CI, Willecke K (2004) Expression pattern of lacZ reporter gene representing connexin36 in transgenic mice. The Journal of comparative neurology 473:511–525.

Ennis M, Zhu M, Heinbockel T, Hayar A (2006) Olfactory nerve-evoked, metabotropic glutamate receptor-mediated synaptic responses in rat olfactory bulb mitral cells. Journal of neurophysiology 95:2233–2241.

Frederick DE, Brown A, Tacopina S, Mehta N, Vujovic M, Brim E, Amina T, Fixsen B, Kay LM (2017) Task-Dependent Behavioral Dynamics Make the Case for Temporal Integration in Multiple Strategies during Odor Processing. The Journal of neuroscience: the official journal of the Society for Neuroscience 37:4416–4426.

Friedrich RW, Wiechert MT (2014) Neuronal circuits and computations: pattern decorrelation in the olfactory bulb. FEBS Lett 588:2504–2513.

Frisch C, De Souza-Silva MA, Sohl G, Guldenagel M, Willecke K, Huston JP, Dere E (2005) Stimulus complexity dependent memory impairment and changes in motor performance after deletion of the neuronal gap junction protein connexin36 in mice. Behavioural brain research 157:177–185.

Galan RF, Fourcaud-Trocme N, Ermentrout GB, Urban NN (2006) Correlation-induced synchronization of oscillations in olfactory bulb neurons. The Journal of neuroscience: the official journal of the Society for Neuroscience 26:3646–3655.

Gire DH, Schoppa NE (2009) Control of on/off glomerular signaling by a local GABAergic microcircuit in the olfactory bulb. The Journal of neuroscience: the official journal of the Society for Neuroscience 29:13454–13464.

Gire DH, Zak JD, Bourne JN, Goodson NB, Schoppa NE (2019) Balancing Extrasynaptic Excitation and Synaptic Inhibition within Olfactory Bulb Glomeruli. eNeuro 6.

Gire DH, Franks KM, Zak JD, Tanaka KF, Whitesell JD, Mulligan AA, Hen R, Schoppa NE (2012) Mitral cells in the olfactory bulb are mainly excited through a multistep signaling path. The Journal of neuroscience: the official journal of the Society for Neuroscience 32:2964–2975.

Gschwend O, Beroud J, Vincis R, Rodriguez I, Carleton A (2016) Dense encoding of natural odorants by ensembles of sparsely activated neurons in the olfactory bulb. Scientific reports 6:36514.

Hayar A, Shipley MT, Ennis M (2005) Olfactory bulb external tufted cells are synchronized by multiple intraglomerular mechanisms. The Journal of neuroscience: the official journal of the Society for Neuroscience 25:8197–8208.

Hayar A, Karnup S, Shipley MT, Ennis M (2004) Olfactory bulb glomeruli: external tufted cells intrinsically burst at theta frequency and are entrained by patterned olfactory input. The Journal of neuroscience: the official journal of the Society for Neuroscience 24:1190–1199.

Heinbockel T, Heyward P, Conquet F, Ennis M (2004) Regulation of main olfactory bulb mitral cell excitability by metabotropic glutamate receptor mGluR1. Journal of neurophysiology 92:3085–3096.

Imamura F, Ito A, LaFever BJ (2020) Subpopulations of Projection Neurons in the Olfactory Bulb. Frontiers in neural circuits 14:561822.

Isaacson JS (1999) Glutamate spillover mediates excitatory transmission in the rat olfactory bulb. Neuron 23:377–384.

Jones S, Zylberberg J, Schoppa N (2020) Cellular and Synaptic Mechanisms That Differentiate Mitral Cells and Superficial Tufted Cells Into Parallel Output Channels in the Olfactory Bulb. Frontiers in cellular neuroscience 14:614377.

Kashiwadani H, Sasaki YF, Uchida N, Mori K (1999) Synchronized oscillatory discharges of mitral/tufted cells with different molecular receptive ranges in the rabbit olfactory bulb. Journal of neurophysiology 82:1786–1792.

Kay LM (2014) Circuit oscillations in odor perception and memory. Progress in brain research 208:223–251.

Kosaka T, Kosaka K (2005) Intraglomerular dendritic link connected by gap junctions and chemical synapses in the mouse main olfactory bulb: electron microscopic serial section analyses. Neuroscience 131:611–625.

Kosaka T, Deans MR, Paul DL, Kosaka K (2005) Neuronal gap junctions in the mouse main olfactory bulb: morphological analyses on transgenic mice. Neuroscience 134:757–769.

Lagier S, Carleton A, Lledo PM (2004) Interplay between local GABAergic interneurons and relay neurons generates gamma oscillations in the rat olfactory bulb. The Journal of neuroscience: the official journal of the Society for Neuroscience 24:4382–4392.

Lepousez G, Lledo PM (2013) Odor discrimination requires proper olfactory fast oscillations in awake mice. Neuron 80:1010–1024.

Lepousez G, Mouret A, Loudes C, Epelbaum J, Viollet C (2010) Somatostatin contributes to in vivo gamma oscillation modulation and odor discrimination in the olfactory bulb. The Journal of neuroscience: the official journal of the Society for Neuroscience 30:870–875.

Logothetis NK, Schall JD (1989) Neuronal correlates of subjective visual perception. Science 245:761–763.

Losacco J, Ramirez-Gordillo D, Gilmer J, Restrepo D (2020) Learning improves decoding of odor identity with phase-referenced oscillations in the olfactory bulb. eLife 9.

Maher BJ, McGinley MJ, Westbrook GL (2009) Experience-dependent maturation of the glomerular microcircuit. Proceedings of the National Academy of Sciences of the United States of America 106:16865–16870.

Mori K, Sakano H (2021) Olfactory Circuitry and Behavioral Decisions. Annu Rev Physiol 83:231–256.

Najac M, De Saint Jan D, Reguero L, Grandes P, Charpak S (2011) Monosynaptic and polysynaptic feed-forward inputs to mitral cells from olfactory sensory neurons. The Journal of neuroscience: the official journal of the Society for Neuroscience 31:8722–8729.

Pouille F, McTavish TS, Hunter LE, Restrepo D, Schoppa NE (2017) Intraglomerular gap junctions enhance interglomerular synchrony in a sparsely connected olfactory bulb network. The Journal of physiology 595:5965–5986.

Rall W, Shepherd GM (1968) Theoretical reconstruction of field potentials and dendrodendritic synaptic interactions in olfactory bulb. Journal of neurophysiology 31:884–915.

Rash JE, Davidson KG, Kamasawa N, Yasumura T, Kamasawa M, Zhang C, Michaels R, Restrepo D, Ottersen OP, Olson CO, Nagy JI (2005) Ultrastructural localization of connexins (Cx36, Cx43, Cx45), glutamate receptors and aquaporin-4 in rodent olfactory mucosa, olfactory nerve and olfactory bulb. J Neurocytol 34:307–341.

Rinberg D, Koulakov A, Gelperin A (2006) Speed-accuracy tradeoff in olfaction. Neuron 51:351–358.

Schoppa NE (2006a) AMPA/kainate receptors drive rapid output and precise synchrony in olfactory bulb granule cells. The Journal of neuroscience: the official journal of the Society for Neuroscience 26:12996–13006.

Schoppa NE (2006b) Synchronization of olfactory bulb mitral cells by precisely timed inhibitory inputs. Neuron 49:271–283.

Schoppa NE, Westbrook GL (2001) Glomerulus-specific synchronization of mitral cells in the olfactory bulb. Neuron 31:639–651.

Schoppa NE, Westbrook GL (2002) AMPA autoreceptors drive correlated spiking in olfactory bulb glomeruli. Nature neuroscience 5:1194–1202.

Shepherd GM, Chen, W. R., Greer, C. A. (2004) Olfactory Bulb. In: The Synaptic Organization of the Brain (Shepherd GM, ed). New York, New York: Oxford University Press.

Stopfer M, Bhagavan S, Smith BH, Laurent G (1997) Impaired odour discrimination on desynchronization of odour-encoding neural assemblies. Nature 390:70–74.

Tan J, Savigner A, Ma M, Luo M (2010) Odor information processing by the olfactory bulb analyzed in gene-targeted mice. Neuron 65:912–926.

Uchida N, Mainen ZF (2003) Speed and accuracy of olfactory discrimination in the rat. Nature neuroscience 6:1224–1229.

Urban NN, Sakmann B (2002) Reciprocal intraglomerular excitation and intra- and interglomerular lateral inhibition between mouse olfactory bulb mitral cells. The Journal of physiology 542:355–367.

Vaaga CE, Westbrook GL (2016) Parallel processing of afferent olfactory sensory information. The Journal of physiology 594:6715–6732.

Vaaga CE, Westbrook GL (2017) Distinct temporal filters in mitral cells and external tufted cells of the olfactory bulb. The Journal of physiology 595:6349–6362.

Yokoi M, Mori K, Nakanishi S (1995) Refinement of odor molecule tuning by dendrodendritic synaptic inhibition in the olfactory bulb. Proceedings of the National Academy of Sciences of the United States of America 92:3371–3375.

Yuan Q, Knopfel T (2006) Olfactory nerve stimulation-evoked mGluR1 slow potentials, oscillations, and calcium signaling in mouse olfactory bulb mitral cells. Journal of neurophysiology 95:3097–3104.

Zlomuzica A, Viggiano D, Degen J, Binder S, Ruocco LA, Sadile AG, Willecke K, Huston JP, Dere E (2012) Behavioral alterations and changes in Ca/calmodulin kinase II levels in the striatum of connexin36 deficient mice. Behavioural brain research 226:293–300.

